# PD-1 inhibition drives CD8^+^ T cell recruitment and T cell-dependent myeloid remodeling in responsive tumors

**DOI:** 10.64898/2026.11.17.767318

**Authors:** Oluwatoyosi Adewunmi, Autumn M. Sunderland, Alec J. Solis, Arielle G. Dessens, Aloukick K. Singh, Matthew M. Gubin, Stephanie S. Watowich, Jason M. Schenkel, Kristen E. Pauken

## Abstract

While leukocyte recruitment is central to anti-tumor immunity, how recruitment changes over time and following perturbation with immunotherapy is poorly understood because of a lack of tools to identify recruited leukocytes. We used intravenous antibody labeling to characterize recruited leukocytes in murine tumors and evaluated how programmed cell death protein-1 (PD-1) immune checkpoint inhibitors (ICI) impacted recruitment. Early PD-1 ICI administration, which induced robust tumor control, increased the density of recently recruited CD8^+^ T cells, and these T cells exhibited a less-exhausted phenotype compared to unlabeled intratumoral CD8^+^ T cells. Early PD-1 ICI also caused a shift in myeloid populations to a proinflammatory iNOS^+^ phenotype. Delaying PD-1 ICI resulted in ineffective tumor control, which was associated with reduced leukocyte entry per unit volume, less granzyme B upregulation in CD8^+^ T cells, and less iNOS induction in myeloid cells. Use of FTY720 to prevent lymphocyte egress from lymph nodes abolished the protective effects of early PD-1 ICI and reduced CD8^+^ T cell recruitment. While FTY720 did not impact myeloid cell recruitment, it did reduce myeloid iNOS induction. Lastly, PD-1 deletion in CD8^+^ T cells was sufficient to increase CD8^+^ T cell recruitment to tumors and iNOS expression in myeloid cells. These data suggest that CD8^+^ T cell recruitment both increases the quantity and quality of CD8^+^ T cells in the tumor and drives T cell-driven remodeling of the myeloid compartment. These findings support efforts to develop therapies that promote CD8^+^ T cell recruitment to improve immunotherapy responses.

## Introduction

Immunotherapy agents targeting the programmed cell death protein-1 (PD-1) pathway have revolutionized cancer care, unleashing the patient’s own immune system to eliminate cancer. These inhibitors have shown remarkable successes, driving cures in patients with otherwise terminal disease [1, 2]. However, while a subset of patients achieves durable responses, most fail to experience long-term disease regression [1, 2]. The tumor microenvironment (TME) is hostile to tumor-infiltrating lymphocytes (TILs), with chronic antigen, inflammation, hypoxia, and immunosuppressive pathways contributing to the erosion of TIL functions and resulting in an exhausted T cell transcriptional and epigenetic state that develops over time [3–6]. Similarly, tumors are highly effective at polarizing cells of the innate immune system to be more immunosuppressive, promoting tumor growth [7, 8]. Significant efforts have focused on developing combination strategies to better leverage productive anti-tumor leukocytes and/or diminish the suppressive capacity of immunosuppressive leukocytes within the TME [1, 9]. However, getting the right combination of therapies to the right patient at the right time remains a major clinical challenge [1].

Considering the difficulty in reprogramming of T cell exhaustion and/or immunosuppressive leukocyte behavior after extended periods of time in tumors, we reasoned that a strategy to overcome this barrier would be to drive the migration of new leukocytes from the periphery into the TME. To initiate productive immune responses, many innate leukocytes and all adaptive leukocytes must migrate from the periphery into inflamed tissue sites, including tumors [10]. Upon entry into tumors, leukocytes undergo dramatic phenotypic and functional shifts as they adapt from the blood microenvironment to the TME [10]. We reasoned that recently recruited leukocytes would be less differentiated and/or more plastic than their tumor-residing counterparts, and therefore more capable of being productively leveraged with immunotherapy. To date, however, testing how different types of cancer therapy impact the recruitment of diverse leukocyte subsets has been challenging due to a lack of tools to identify recently recruited leukocytes in tissues. The gold standard for tracking leukocyte recruitment in the field of tissue resident memory has been parabiosis, which involves the surgical conjoining of mice, causing them to share a vascular system. However, parabiosis requires major surgery and it takes 1-2 weeks for full vascularization and equilibration of circulating immune cells between mice, favoring long-term studies of trafficking rather than short-term [11]. An alternative approach is to use photoactivatable dyes encoded in the germline. However, this requires extensive breeding, the penetration of light into tissue to activate the dyes can be limited, and only a single labeling event can be studied [12, 13]. To circumvent these technical limitations, we used an intravenous (IV) antibody-based labeling method to investigate how PD-1 inhibitors modulate migration to murine tumors. This approach involves injecting a fluorescently labeled antibody against the pan-leukocyte marker CD45 into the vasculature of mice [14–16]. This method is adapted from previous studies that performed short-term labeling (3-5 minutes) to distinguish cells in the vasculature from cells in the tissue parenchyma [17, 18]. Newer work has established that this methodology can also be used to identify leukocytes that have been recruited from the blood into tissues in mice [14–16] as well as non-human primates [19, 20].

In this study, we used this novel IV labeling methodology to examine the dynamics of leukocyte recruitment into transplantable tumors at steady state and following PD-1 ICI. By delivering IV antibodies conjugated to different fluorochromes at 24 hours and 3 minutes, we were able to profile how recently recruited (24-hr IV^+^ 3-min IV^−^) leukocytes differed in phenotype and number compared to their unlabeled counterparts (IV^−^), while excluding leukocytes that were still actively in the vasculature (3-min IV^+^). In MC38 tumors we found that CD8^+^ T cell and myeloid cell subsets were continually recruited to tumors over time as tumors progressed; however, the overall number of recently recruited leukocytes per unit volume decreased at later stages of disease. Delivery of PD-1 ICI at earlier stages of disease resulted in an increased density of recently recruited CD8^+^ T cells, and these T cells had more granzyme B expression than the control-treated counterparts. Delivery of PD-1 ICI at later stages of disease resulted in ineffective tumor control and reduced T cell recruitment. Interestingly, the majority of leukocytes within the 24-hr infiltrate were myeloid cells, not T cells, even with PD-1 ICI. While the total number of recently recruited 24hr IV^+^ F4/80^+^ MHCII^+^ and Ly6C^+^ myeloid cells did not change following PD-1 ICI, we observed an increase in the frequency of both myeloid populations that expressed the proinflammatory marker iNOS at earlier stages of disease but not later stages of disease. Blunting lymphocyte migration using the drug FTY720 abolished the protective effects of PD-1 ICI on tumor growth, CD8^+^ T cell recruitment, and myeloid cell iNOS induction, suggesting that lymphocyte recruitment was critical for these events. Lastly, restricting PD-1 loss to the CD8^+^ T cell compartment was sufficient to drive increased CD8^+^ T cell recruitment and myeloid cell induction of iNOS, highlighting the importance of PD-1 inhibitors acting on CD8^+^ T cells to drive both T cell recruitment to the tumor and myeloid cell remodeling to a more proinflammatory phenotype. Collectively, these findings support a model where successful PD-1 ICI is associated with increased recruitment of proinflammatory, anti-tumor innate and adaptive leukocytes to tumors, while unsuccessful PD-1 ICI fails to adequately mobilize these cells. These results provide a rationale for targeting leukocyte migration in future studies to improve immunotherapy responses in patients, as well as leveraging monitoring of peripheral leukocytes to assess PD-1 response.

## Results

### IV labeling reveals dynamic recruitment of innate and adaptive leukocytes into tumors

To date, assessing leukocyte entry into tumors and characterizing how recently recruited leukocytes differ from non-recently recruited leukocytes has been challenging due to a lack of tools to readily identify recently recruited leukocytes in native, unperturbed tissues. Here, we used a recently described IV antibody labeling approach [14, 15] to quantify leukocyte entry rate and phenotype in the TME. Using subcutaneous (s.c) tumor models, we first examined the recruitment dynamics of relevant cell populations over time at homeostasis. We started with s.c. implantation of MC38 colon adenocarcinoma cells, since this cell line is highly responsive to PD-1 ICI at early stages of disease [21–23] (**Fig. 1**). We also performed parallel analyses in transplantable melanoma models (B16-OVA, YUMM3.3) which show more of a moderate response to PD-1 ICI [24–27] (**Fig. S1**). We first characterized the dynamics of leukocyte recruitment into these tumors over time, performing serial harvests over tumor progression (**Fig. 1A**). Mice were injected IV with anti-CD45.2 antibodies (clone 104, which is optimal for longer term *in vivo* labeling) at 24 hours and 3 minutes prior to each harvest as previously described [14, 15] to identify recently recruited leukocytes (defined as 24-hr IV^+^ 3-min IV^−^) as well as leukocytes that were actively in the vasculature (3-min IV^+^) (**Fig. 1B**). The anti-CD45.2 given at 24 hours and 3 minutes were conjugated to different fluorophores so these labels could be readily distinguished (**Fig. 1B**). We consistently observed a small population of leukocytes that bind the 3-min IV label indicating that they were actively in the tumor vasculature at the time of harvest (**Fig. 1B**), and these cells were excluded from downstream analyses. The leukocytes that were 24-hr IV^+^ 3-min IV^−^ were in the blood at 24 hours when the rapid labeling event occurred, but were in the tumor at time of harvest (**Fig. 1B**). The leukocytes that bind neither IV antibody (defined as 24-hr IV^−^ 3-min IV^−^) have likely been in the tumor for longer than 24 hours (**Fig. 1B**). However, since the active labeling period at 24 hours is short (∼45 minutes) [14], it is possible that there may be some recently recruited leukocytes in the unlabeled fraction in tumors.

**Figure 1:**
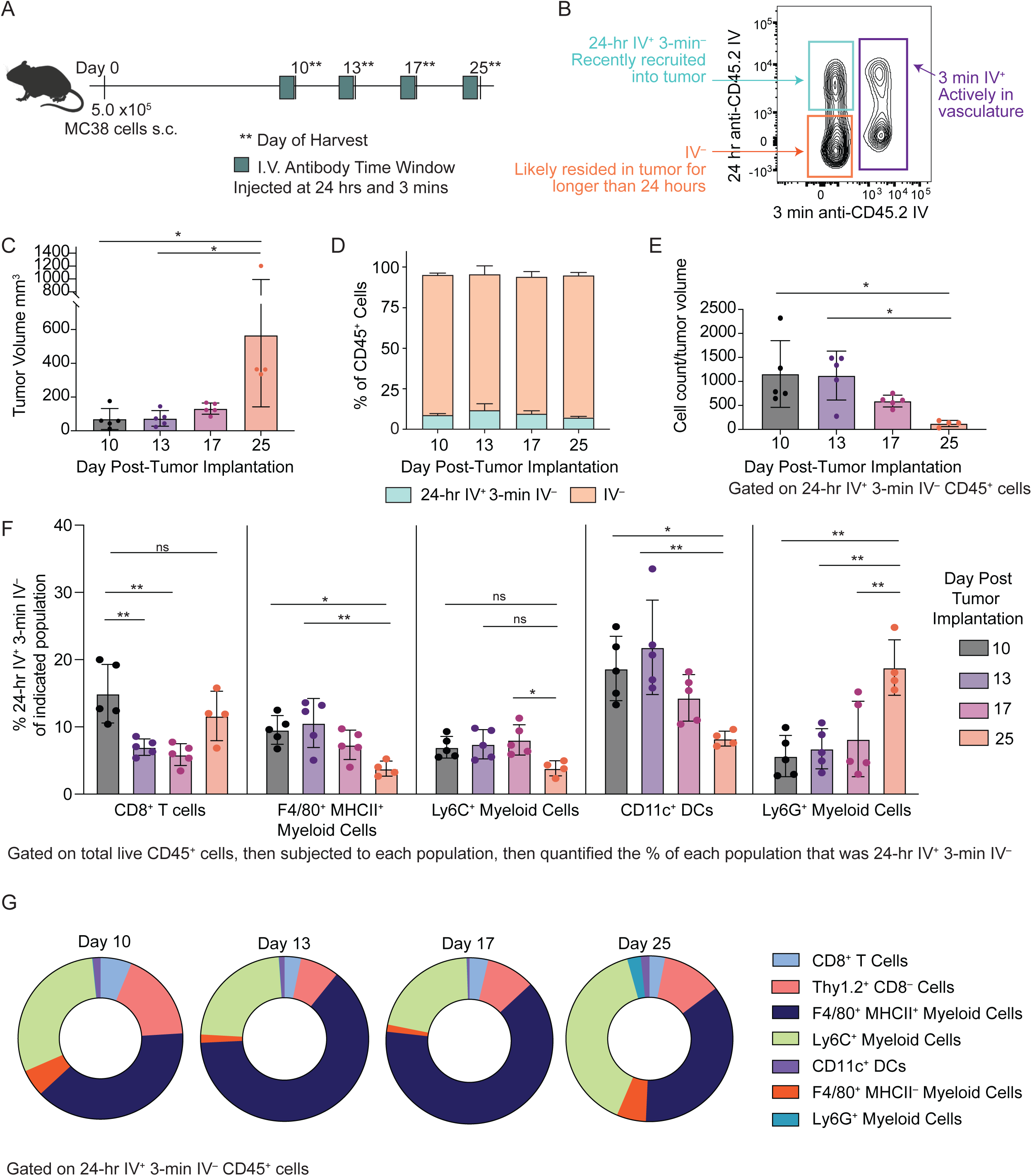
IV antibody labeling shows dynamic recruitment of diverse innate and adaptive leukocyte subsets to MC38 tumors over time. **A.** Schematic of experimental design. Female C57BL/6 mice were injected subcutaneously with 500,000 MC38 cells and randomized into day 10, 13, 17, and 25 harvest cohorts. 24 hours and 3 minutes prior to harvest, mice were injected with anti-CD45.2 antibodies conjugated to different fluorophores IV to label cells in the vasculature. **B.** Representative flow plot of CD45^+^ cells from a MC38 tumor (day 10) showing 24-hr vs. 3-min IV labeling with anti-CD45.2 antibody. Cells were gated on live, CD45^+^. Cells are defined as being “recently recruited” (24-hr IV^+^ 3-min IV^−^, top left quadrant) or “actively in the vasculature” (3-min IV^+^, right quadrant). Cells that lack both IV antibodies (IV^−^, bottom left quadrant) were not present in the blood at either 24 hours or 3 minutes. **C.** Tumor volume at time of tumor harvest (n = 4-5 mice per group). Representative of two independent experiments. **D-G** show quantification of flow cytometry data of leukocytes from MC38 tumors. **D.** Percentage of total CD45^+^ cells that were 24-hr IV^+^ 3-min IV^−^ vs. total CD45^+^ cells that were IV^−^. **E.** Total cell count of recently recruited CD45^+^ immune cells (24-hr IV^+^ 3-min IV^−^) normalized to tumor volume. **F.** Frequency of each indicated immune cell lineage that was recently recruited (24-hr IV^+^ 3-min IV^−^) over time. All lineages are gated on live, CD45^+^. Individual lineages further defined as: CD8^+^ <u>T cells</u>: Thy1.2^+^ CD8^+^; F4/80^+^ MHCII^+^ myeloid cells: F4/80^+^, Ly6G^−^, Ly6C^−^, MHCII^+^, CD11b^+^; Ly6C^+^ myeloid cells: F4/80^+^, Ly6G^−^, Ly6C^+^; CD11c^+^ dendritic cells (DCs): F4/80^−^, Ly6G^−^, CD11c^+^, MHCII^+^; Ly6G^+^ myeloid cells: F4/80^−^, Ly6G^+^. Gating strategy shown in **Fig. S1C**. **G**. Composition of recently recruited (24-hr IV^+^ 3-min IV^−^) immune cell population over time. Gated on total CD45^+^ and 24-hr IV^+^ 3-min IV^−^ first, and then applied the lineage-specific gating shown in **Fig. 1F** to assess the composition of the 24-hr IV^+^ infiltrate, with the addition of F4/80^−^ MHCII^−^ myeloid cells defined as F4/80^+^, Ly6G^−^, Ly6C^−^, MHCII^−^, CD11b^+^ and CD8^−^ T cells defined as Thy1.2^+^ CD8^−^. Data shown in panels **Fig. 1C-1G** are representative of two independent experiments with 3-5 mice per group per experiment. Statistics calculated using One-way ANOVA with multiple comparisons tests. In the Figure, ns indicates p>0.05, * indicates p<0.05, and ** indicates p<0.01.

As expected, MC38 tumors increased in size over time (**Fig. 1C**). At every time point examined, we observed a consistent fraction of the total leukocytic infiltrate was labeled with the 24-hour IV antibody, albeit small compared to the IV⁻ fraction (mean ± SD of 24-hr IV⁺ 3-min IV⁻ cells within total CD45⁺ cells: day 10 = 12.38% ± 1.0; day 13 = 19.54% ± 6.5; day 17 = 18.03% ± 2.8; day 25 = 11.24% ± 1.0) (**Fig. 1D**). However, when normalized to tumor volume, the density of 24-hour IV^+^ 3-min IV^−^ leukocytes decreased as tumors progressed (**Fig. 1E**), consistent with immune evasion at later stages of disease. We observed similar recruitment dynamics to s.c. MC38 in the s.c melanoma model B16-OVA (**Fig. S1A**). In YUMM3.3 s.c. melanoma tumors, we observed a similar trend to MC38 and B16-OVA in that the frequency of recently recruited leukocytes of total CD45^+^ was consistent across time points (**Fig. S1B**). However, in YUMM3.3, we noted that the peak of recruitment in terms of cell number per tumor volume occurred later than MC38 and B16-OVA (**Fig. S1B**). These data support a model where leukocyte recruitment to tumors continually occurs over time, though as tumors grow, the quantity of recently recruited leukocytes per unit volume decreases.

We next examined the entry rate of different leukocyte subsets in the tumor, including CD8^+^ T cells, F4/80^+^ MHCII^+^ Ly6C^−^ myeloid cells (likely macrophages), Ly6C^+^ myeloid cells (likely monocytes), CD11c^+^ MHCII^+^ dendritic cells (DCs), and Ly6G^+^ myeloid cells (likely neutrophils) (gating strategy shown in **Fig. S1C**). To do this, we gated first on each of these subpopulations, and then examined the frequency of recently recruited (24-hour IV^+^ 3-min IV^−^) cells within each population. Although there was variability across the different leukocyte subtypes, the overall fraction of recently recruited leukocytes within each major population was relatively low in MC38 tumors (**Fig. 1F**). For CD8^+^ T cells, the frequency of recently recruited cells within the total population was highest at day 10 and decreased at subsequent time points (**Fig. 1F**). The frequency of the F4/80^+^ MHCII^+^ myeloid and Ly6C^+^ myeloid populations that had recently recruited into the tumor was relatively constant over time, with around 10% of F4/80^+^ MHCII^+^ myeloid and 7% of Ly6C^+^ myeloid cells having recently entered the tumor, though a drop at day 25 was noted (3.8 ± 1.1 of F4/80^+^ MHCII^+^ myeloid and 3.8% ± 1.1 of Ly6C^+^ myeloid cells). The DC population showed 15-20% of cells being recently recruited into the tumor (**Fig. 1F**). Very few Ly6G^+^ myeloid cells were observed in MC38 tumors, and their entry rate was ∼6-8% until the final time point (18.8 ± 4.1) (**Fig. 1F**). In B16-OVA melanoma (**Fig. S1D**) and YUMM3.3 melanoma (**Fig. S1E**), recruitment was relatively consistent across time points. However, these tumors contained a larger population of Ly6G^+^ myeloid cells compared with MC38, and the entry rates of the F4/80^+^ MHCII^+^ myeloid, Ly6C^+^ myeloid, and Ly6G^+^ myeloid populations were generally higher than in MC38 (**Fig. 1F, S1D, S1E**).

Lastly, we gated on the 24-hour IV^+^ 3-min IV^−^ population first and then quantified the distribution of different leukocyte subsets within that population. This analysis enabled us to determine which immune cells were actively entering into the tumor, regardless of the size of their corresponding intratumoral population. When we characterized the 24-hour IV^+^ 3-min IV^−^ fraction, CD8^+^ T cells were the minority of leukocytes entering the tumor at steady state at all time points examined (day 10, 6.8% ± 2.2; day 13, 3.3% ± 1.3; day 17, 3.9% ± 1.2; day 25; 2.9%±1.1) (**Fig. 1G**). Rather, the majority of the new leukocytes entering tumors were either Ly6C^+^ myeloid cells or F4/80^+^ MHCII^+^ myeloid cells (**Fig. 1G**). We observed a small population of F4/80^+^ MHCII^−^ Ly6C^−^ myeloid cells at each time point (∼5% or less). Very few Ly6G^+^ myeloid cells (likely neutrophils) were observed at any time point until day 25 in MC38 (2.5% ± 1.5). Collectively, these data show that leukocyte entry into tumors is continually occurring over time, although the number of newly recruited cells decreases relative to tumor size over time. These findings also reveal continuous leukocyte entry into tumors that is dominated by myeloid cells, and relatively limited recruitment of CD8⁺ T cells over time.

### Recently recruited CD8^+^ T cells and myeloid cells display phenotypic differences compared to IV^−^ cells

We next characterized the phenotype of the recently recruited (24-hr IV^+^ 3-min IV^−^) leukocyte infiltrate compared to IV^−^ leukocytes. Since time in the TME is thought to be a driver of both CD8^+^ T cell exhaustion and myeloid skewing to an immunosuppressive phenotype [10, 28–30], we speculated that recently recruited leukocytes would have a less differentiated phenotype (e.g., CD8^+^ T cells would be less terminally exhausted, and myeloid cells would be less immunosuppressive). At steady state, 24-hr IV^+^ 3-min IV^−^ CD8^+^ TILs expressed lower levels of individual inhibitory receptors, including PD-1 (**Fig. 2A**) as well as TIM-3, LAG-3, and TIGIT (**Fig. S2A-S2C**). Previous reports have shown that co-expression of multiple inhibitory receptors correlates with the degree of exhaustion in both chronic viral infection and cancer [31, 32]. We observed that 24-hr IV^+^ 3-min IV^−^ CD8^+^ TILs showed lower co-expression of inhibitory receptors (PD-1, CD160, TIGIT, TIM-3, and LAG-3) than their IV^−^ counterparts (**Fig. 2B**, **2C**), consistent with recently recruited T cells having a less exhausted phenotype. These findings are concordant with prior reports using photoactivatable alleles encoded in the germline, which showed that T cells adopt an exhausted phenotype after being retained in the tumor for 72 hours or longer [33]. Across all time points examined, 24-hr IV^+^ 3-min IV^−^ CD8^+^ TILs had lower levels of inhibitory receptor co-expression while the IV^−^ populations had higher levels of inhibitory receptor co-expression (**Fig. 2B**), suggesting that recently recruited populations display a less exhausted phenotype over time. We also noted lower expression of the cytotoxic effector marker Granzyme B on 24-hr IV^+^ 3-min IV^−^ CD8^+^ TILs compared to their IV^−^ counterparts, which reached significance at the day 10, 13, and 17 time points, consistent with these T cells being less differentiated (**Fig. 2D**). This finding is consistent with previous work showing that upon tumor entry, time is required for full granzyme B upregulation on CD8^+^ T cells [34]. These data suggest that recently recruited CD8^+^ T cells could be a source of less-exhausted cells that could be leveraged by immunotherapy to drive sustained anti-tumor activity throughout tumor progression.

**Figure 2:**
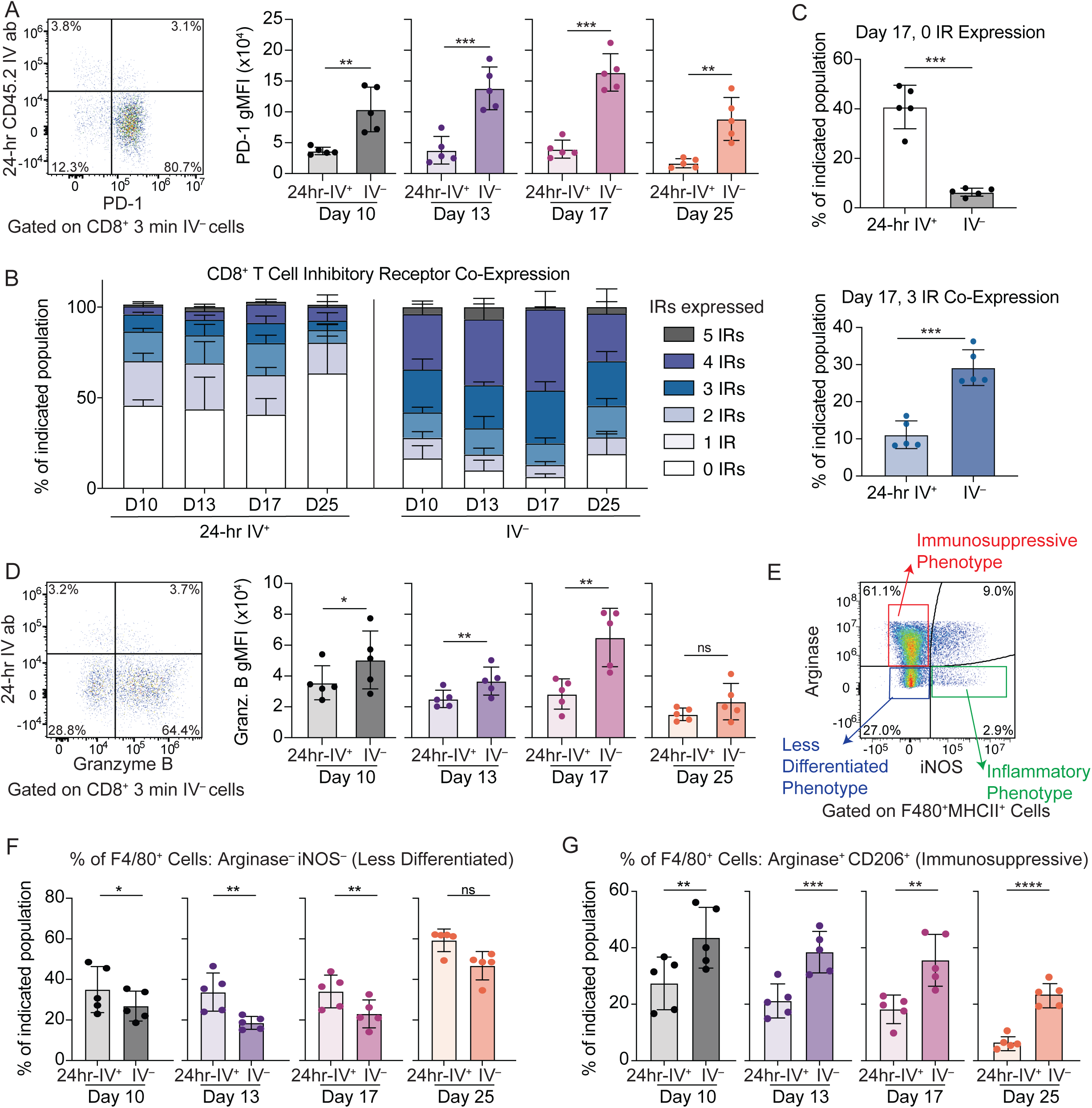
Recently recruited CD8^+^ T cells and F4/80^+^ MHCII^+^ myeloid cells show phenotypic differences compared to their unlabeled counterparts. Phenotypic analyses performed on the mice shown in **Fig. 1**. WT female C57BL/6 mice were injected subcutaneously with 500,000 MC38 cells and harvested on days 10, 13, 17, and 25. 24 hours and 3 minutes prior to harvest, mice were injected with anti-CD45.2 antibodies conjugated to different fluorophores IV. In A – D, cells are gated on live, CD45^+^, Thy1.2^+^, CD8^+^, 3-min IV^−^ to show comparisons of 24-hr IV^+^ vs. IV^−^ CD8^+^ T cells within the MC38 tumor tissue. “24-hr IV^+^” indicates “24-hr IV^+^ 3-min IV^−^”, while “IV^−^” indicates “24-hr IV^−^ 3-min IV^−^”. A. (Left) Representative flow plot showing PD-1 vs. the 24-hr anti-CD45.2 label on CD8^+^ T cells in MC38 tumors (day 17). (Right) Quantification of PD-1 expression using geometric median fluorescent intensity (gMFI) in the indicated CD8^+^ TIL populations at different time points post tumor cell implantation. B. Percentage of the indicated CD8^+^ T cell populations that express or co-express the following inhibitory receptors quantified using flow cytometry. Included in the analysis are PD-1, CD160, TIM-3, TIGIT, and LAG-3. Number of inhibitory receptors expressed/co-expressed determined using combination gating (Boolean gating) in FlowJo software. C. (Top) Quantification of 24-hr IV^+^ 3-min IV^−^ vs. IV^−^ CD8^+^ T cells that express 0 inhibitory receptors at day 17 vs. (bottom) quantification of 24-hr IV^+^ 3-min IV^−^ vs. IV^−^ CD8^+^ T cells that co-express 3 inhibitory receptors at day 17. D. (Left) Representative flow plot showing granzyme B vs. the 24-hr anti-CD45.2 label on CD8^+^ T cells in MC38 tumors (day 17). (Right) Quantification of granzyme B expression in the indicated CD8^+^ TIL populations using gMFI at different time points. E. Representative flow plot showing arginase vs. iNOS expression in F4/80^+^ MHCII^+^ myeloid cells at day 10. F. Frequency of indicated populations of F4/80^+^ MHCII^+^ myeloid cells that were arginase^−^ and iNOS^−^ at different time points. G. Frequency of indicated populations of F4/80^+^ MHCII^+^ myeloid cells that were arginase^+^ CD206^+^ at different time points. In **Fig. 2E – 2G**, F4/80^+^ MHCII^+^ myeloid are defined using the gating strategy F4/80^+^, Ly6G^−^, Ly6C^−^, MHCII^+^, CD11b^+^. Data shown in **Fig. 2A – 2G** are from one experiment with n = 5 mice per group. The data are representative of two independent experiments with 4-5 mice per group per experiment. Statistics calculated using a paired Student’s t test at each individual time point. In the Figure, ns indicates p>0.05, * indicates p<0.05, ** indicates p<0.01, *** indicates p<0.001, and **** indicates p<0.0001.

Next, we characterized key populations of myeloid cells within the recently recruited fraction over time. We used a combination of iNOS and arginase expression to characterize myeloid cells that were less differentiated (defined as iNOS^−^ arginase^−^), more proinflammatory (defined as iNOS^+^ arginase^−^), and more immunosuppressive (defined as iNOS^−^ arginase^+^) (**Fig. 2E**). Within the F4/80^+^ MHCII^+^ myeloid population, we further sub-gated arginase^+^ cells that also expressed CD206, a marker associated with a more immunosuppressive phenotype [35, 36]. Within the F4/80^+^ MHCII^+^ myeloid population, the frequency of less differentiated cells (iNOS^−^ arginase^−^) was higher in the 24-hr IV^+^ 3-min IV^−^ population compared to the IV^−^ population at all time points except day 25, where the trend was still noted but not significant (**Fig. 2F**). Conversely, there was a greater population of immunosuppressive iNOS^−^ arginase^+^ CD206^+^ cells in the IV^−^ F4/80^+^ MHCII^+^ myeloid cell compartment compared to 24-hr IV^+^ 3-min IV^−^ F4/80^+^ MHCII^+^ myeloid cells. (**Fig. 2G**). Interestingly, there were little to no differences in terms of iNOS and arginase expression between the 24-hr IV^+^ 3-min IV^−^ and IV^−^ populations within the Ly6C^+^ myeloid subset (**Fig. S2D, S2E**), which is likely comprised of inflammatory monocytes. This observation was consistent with the model that monocytes rapidly differentiate into other populations (including macrophages, monocyte-derived MDSCs, and/or monocyte-derived DCs) upon entering into the tumor [16, 37], so there was likely less of a time difference between the 24-hr IV^+^ 3-min IV^−^ and IV^−^ Ly6C^+^ myeloid cells than the 24-hr IV^+^ 3-min IV^−^ and IV^−^ F4/80^+^ MHCII^+^ myeloid cells.

Collectively, these findings suggest that there are qualitative differences between recently recruited leukocytes and IV^−^ leukocytes. These data support a model where leukocytes generally enter into the tumor in a less differentiated state, and then further differentiate upon entry into the TME [16, 33]. These differences were more pronounced for CD8^+^ T cells and F4/80^+^ MHCII^+^ myeloid cells, and less pronounced for Ly6C^+^ myeloid cells.

### PD-1 ICI drives a more substantial increase in recruitment of CD8^+^ T cells at earlier stages of disease than later stages of disease

Previous work in s.c. MC38 has shown that PD-1 ICI is more effective at inducing tumor clearance when given in earlier-stage disease and/or small tumors than later-stage disease and/or bigger tumors [21, 22]. We asked whether the difference in the ability of PD-1 ICI to promote tumor clearance in MC38 at earlier vs. later stages of disease correlated with a differential ability of PD-1 ICI to induce recruitment of CD8^+^ T cells to tumors. We administered PD-1 ICI earlier (one dose at day 7, harvest at day 10) or later (doses at day 18 and day 21, harvest at day 24) post-tumor implantation, allowing us to keep the tumor type constant (**Fig. 3A**). One dose of anti-PD-1 given at day 7 led to a reduction in tumor size at the day 10 harvest, while two doses of anti-PD-1 given starting at day 18 did not result in a significant reduction in tumor size at the day 24 harvest (**Fig. 3B**). IV antibody labeling was performed at 24 hours and 3 minutes prior to harvest (**Fig. 3A**). At day 10, we observed a significant increase in the number of both recently recruited (24-hr IV^+^ 3-min IV^−^) CD45^+^ (**Fig. 3C**) and CD8^+^ T cells (**Fig. 3D**) per unit of tumor volume. On the contrary, we did not observe an increase at later stages of disease with anti-PD-1 (**Fig. 3C**, **3D**). We did not observe a significant difference in the overall entry rate of CD8^+^ T cells (e.g., the frequency of 24-hr IV^+^ 3-min IV^−^ within the CD8^+^ T cell population) at either time point (**Fig. S3A**). Despite the differences in the number of recently recruited CD8^+^ T cells depending on tumor stage, we observed that the recently recruited CD8^+^ T cells showed lower co-expression of multiple inhibitory receptors both during early and late stages of disease, suggesting that the T cells that were being recruited were of a less exhausted phenotype regardless of disease stage (**Fig. 3E-3G**). At early stages of disease, anti-PD-1 treatment resulted in an increase in the number of recently recruited CD8^+^ T cells that expressed the cytotoxic molecule granzyme B (**Fig. 3H**), as well as an increase in the geometric mean fluorescence intensity (gMFI) of granzyme B within the recently recruited population (**Fig. 3I**). Similarly, early anti-PD-1 caused an increase in the number of recently recruited CD8^+^ T cells in the tumor that expressed the proliferative marker Ki-67 (**Fig. 3J**). Conversely, at later stages of disease, there were very few CD8^+^ T cells that expressed granzyme B within the 24-hr IV^+^ 3-min IV^−^ fraction regardless of treatment, although the difference between isotype control and anti-PD-1-treated did reach significance (**Fig. 3H**). However, the gMFI was lower at the day 24 timepoint and not significantly different between treatment groups (**Fig. 3I**). There was also no difference observed in the frequency of recently recruited CD8^+^ T cells that expressed Ki-67 between treatment groups at the later timepoint as well. (**Fig. 3J**). Collectively, these data support a model where PD-1 ICI effectively induces the recruitment of less exhausted CD8^+^ T cells to tumors at early stages of disease when these inhibitors are more effective at clearing the tumor, while PD-1 ICI is less effective at inducing recruitment of CD8^+^ T cells at later stages of disease when these inhibitors are less effective at controlling tumor growth.

**Figure 3:**
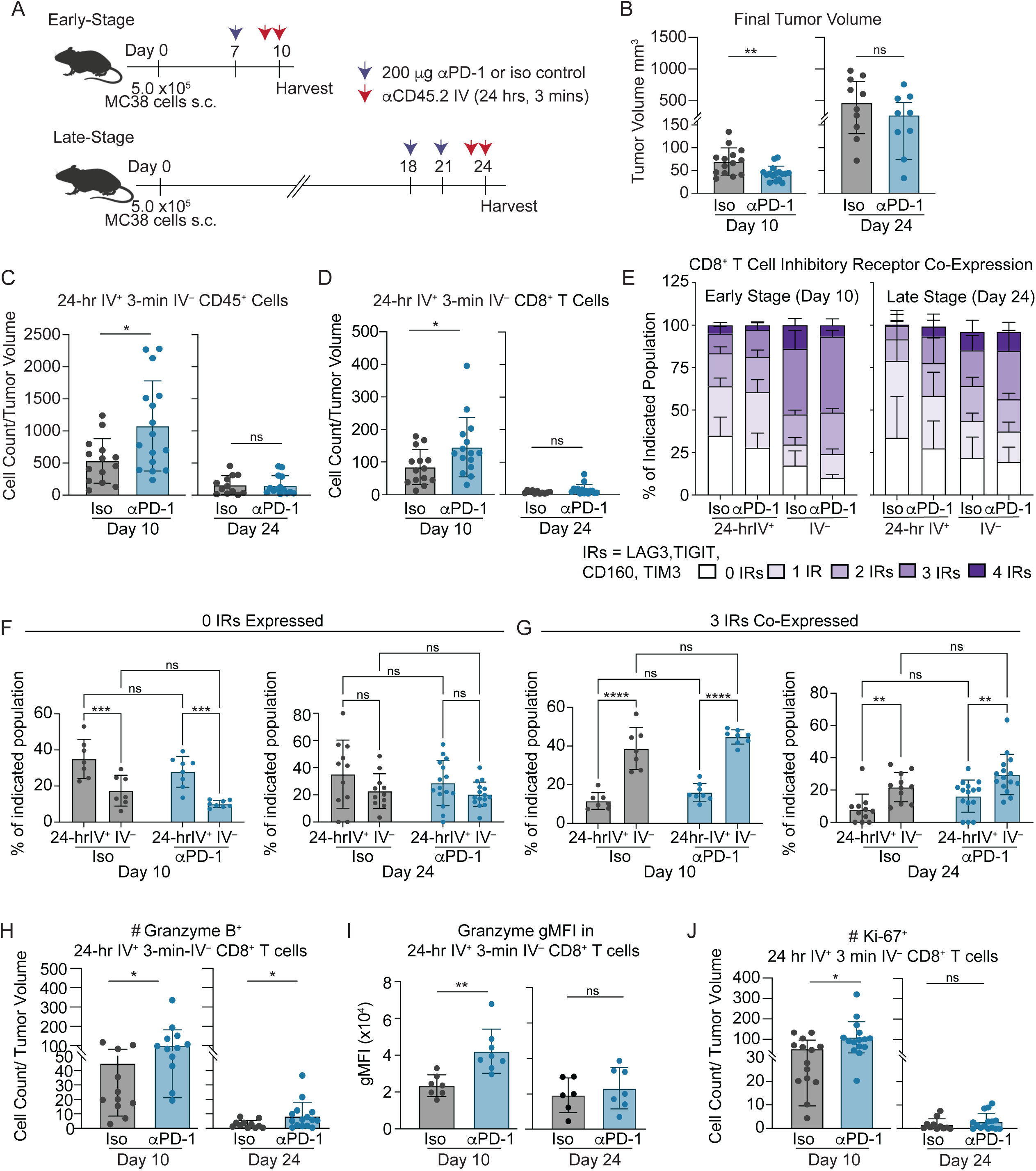
PD-1 ICI drives more substantial recruitment of CD8^+^ T cells into tumors at early stages of disease than later stages of disease. **A.** Experimental design to test how PD-1 ICI given earlier vs. later during MC38 tumor progression impacted leukocyte recruitment. Female C57BL/6 mice were injected subcutaneously with 500,000 MC38 cells and randomized into isotype or anti-PD-1 treatment cohorts. ICI given on day 7 for the early-stage (followed by harvest at day 10) or day 18 and 21 for the late-stage (followed by harvest at day 24). 24 hours and 3 minutes prior to harvest, mice were injected with anti-CD45.2 antibodies conjugated to different fluorophores IV. B. Tumor volume at time of harvest. In C, cells are gated on live, CD45^+^. In D – J, cells are gated on live, CD45^+^, Thy1.2^+^, CD8^+^. C. Total cell count of 24-hr IV^+^ 3-min IV^−^ CD45^+^ cells normalized per unit tumor volume. D. Total cell count of 24-hr IV^+^ 3-min IV^−^ CD8^+^ T cells normalized per unit tumor volume. E. Percentage of the indicated CD8^+^ TIL populations expressing or co-expressing the inhibitory receptors CD160, TIM-3, TIGIT, and/or LAG-3 at the indicated harvest time points. F. Quantification of the indicated CD8^+^ TIL populations expressing 0 inhibitory receptors at the indicated harvest time points. G. Quantification of the indicated CD8^+^ TIL populations that co-express 3 inhibitory receptors labeled at the indicated harvest time points. **H**. Total cell count of granzyme B^+^ recently recruited (24-hr IV^+^ 3-min IV^−^) CD8^+^ TILs normalized to tumor volume at the indicated harvest time points. **I**. Quantification of granzyme B expression using gMFI in recently recruited (24-hr IV^+^ 3-min IV^−^) CD8^+^ TILs at the indicated harvest time points. **J**. Total cell count of Ki-67^+^ recently recruited (24-hr IV^+^ 3-min IV^−^) CD8^+^ TILs normalized to tumor volume at the indicated harvest time points. Data shown in **Fig. 3B – 3J** are a combination of two independent experiments with 7-8 mice per treatment group per experiment and are representative of 2-4 independent experiments. Data shown in **Fig. 3I** are from one experiment (since MFI values cannot be pooled) representative of 2 independent experiments with 7-8 mice per treatment group per experiment. Statistical analyses in **Fig. 3B-3D** and **Fig. 3H- 3J** performed using unpaired Student’s t tests, and analyses in **Fig. 3F** and **3G** performed using 2-way ANOVA with multiple comparisons. In the Figure, ns indicates p>0.05, * indicates p<0.05, ** indicates p<0.01, *** indicates p<0.001, and **** indicates p<0.0001. Throughout the Figure, “24-hr IV^+^” indicates “24-hr IV^+^ 3-min IV^−^”, while “IV^−^” indicates “24-hr IV^−^ 3-min IV^−^”.

In addition to the differences observed within the recently recruited leukocyte populations, we also observed some differences in the IV^−^ compartment (**Fig. S3**). At the earlier time point when PD-1 ICI was more effective, there was a trend towards an increase in total CD45^+^ IV^−^ leukocytes, which did not reach significance (**Fig. S3B**). We did, however, observe an increase in the number of CD8^+^ T cells, granzyme B^+^ CD8^+^ T cells, and Ki-67^+^ CD8^+^ T cells within the IV^−^ fraction at the earlier time point (**Fig. S3C-S3E**), consistent with previous reports using mice with photoactivatable dyes encoded in the germline which showed that both recently recruited and non-recently recruited CD8^+^ T cells show changes following PD-1 inhibition [33]. At the later time point, we did not observe an increase in total CD45^+^ IV^−^ leukocytes (**Fig. S3B**) or IV^−^ Ki-67^+^ CD8^+^ T cells (**Fig. S3E**). We observed slight but significant increases in the number of IV^−^ CD8^+^ T cells and IV^−^ granzyme B^+^ CD8^+^ T cells; however, the differences observed were more minimal in terms of the magnitude of differences at the day 24 time point compared to the day 10 timepoint (**Fig. S3B-S3E**). These findings support a model where PD-1 inhibitors can both induce recruitment of CD8^+^ T cells from the periphery to the tumor as well as reinvigorate tumor-residing populations, consistent with previous work [33]. However, these findings also support a model in which both recruitment and local reinvigoration are less effective when PD-1 ICI is administered at later stages of disease.

### PD-1 blockade remodels myeloid populations in tumors to be more proinflammatory when given at early stages of disease but not later stages of disease

Myeloid cells can have a mixed impact on the ensuing immune response to cancer. Some myeloid cell subsets (including cDC1s and M1-like macrophages) can promote productive anti-tumor immunity, displaying proinflammatory phenotypes. However, many cancer types can hijack myeloid cells to promote tumor growth, supporting the differentiation of these cells into an immunosuppressive phenotype [7, 38, 39]. While some myeloid cells can originate from tissue resident precursors, other myeloid populations are continuously recruited into tissues from the circulation, arising from hematopoietic stem cell progenitors in the bone marrow [40, 41]. In MC38 tumors, we observed that the majority of leukocytes found in the 24-hr IV^+^ 3-min IV^−^ infiltrate were myeloid-derived in both the isotype control-treated and anti-PD-1-treated groups, with the F4/80^+^ MHCII^+^ and Ly6C^+^ myeloid subsets being the most abundant (**Fig. S4A**). Because of this, we next interrogated how PD-1 ICI impacted the myeloid cell compartment. We compared early vs. late stages of disease (**Fig. 3A**), where PD-1 ICI was more or less effective at promoting productive anti-tumor immunity, respectively (**Fig. 3B**). Despite PD-1 inhibitors inducing an increase in the number of total CD45^+^ cells that had recently entered the TME (**Fig. 3C**), we did not observe a significant increase in the overall number of recently recruited F4/80^+^ MHCII^+^ myeloid cells (**Fig. 4A**) or Ly6C^+^ myeloid cells (**Fig. 4B**) when normalizing for tumor volume. However, within the 24-hr IV^+^ 3-min IV^−^ infiltrate, we observed changes to the composition of the F4/80^+^ MHCII^+^ and Ly6C^+^ myeloid populations with PD-1 blockade (**Fig. 4C**, **4D**). There was a significant increase in the iNOS^+^ fractions (both iNOS^+^ arginase^−^ and iNOS^+^ arginase^+^) within both the 24-hr IV^+^ 3-min IV^−^ and IV^−^ fractions at earlier stages of disease (**Fig. 4E-4H, S4B-S4F**), consistent with the notion that PD-1 inhibitors induce an increase in iNOS production by both recently recruited and local myeloid cells in the tumor. Importantly, these changes were far less prominent at the later stages of disease (**Fig. 4C-4H, S4B-S4F**). Collectively, these data suggest that delivery of anti-PD-1 at earlier stages of disease is associated with increased recruitment of proinflammatory myeloid populations as well as iNOS induction in IV^−^ myeloid cells, while delivery of anti-PD-1 at later stages of disease fails to promote the accumulation of proinflammatory myeloid subsets. We also noted a decrease in the immunosuppressive arginase^+^ iNOS^−^ myeloid cell compartments at early stage of disease in both recently recruited and IV^−^ populations but not later stage disease (**Fig. 4C-4D, S4G-S4J**). These data are consistent with the iNOS findings, suggesting PD-1 immunotherapy is promoting a shift to a more proinflammatory phenotype at early stages of disease but not later stages of disease.

**Figure 4:**
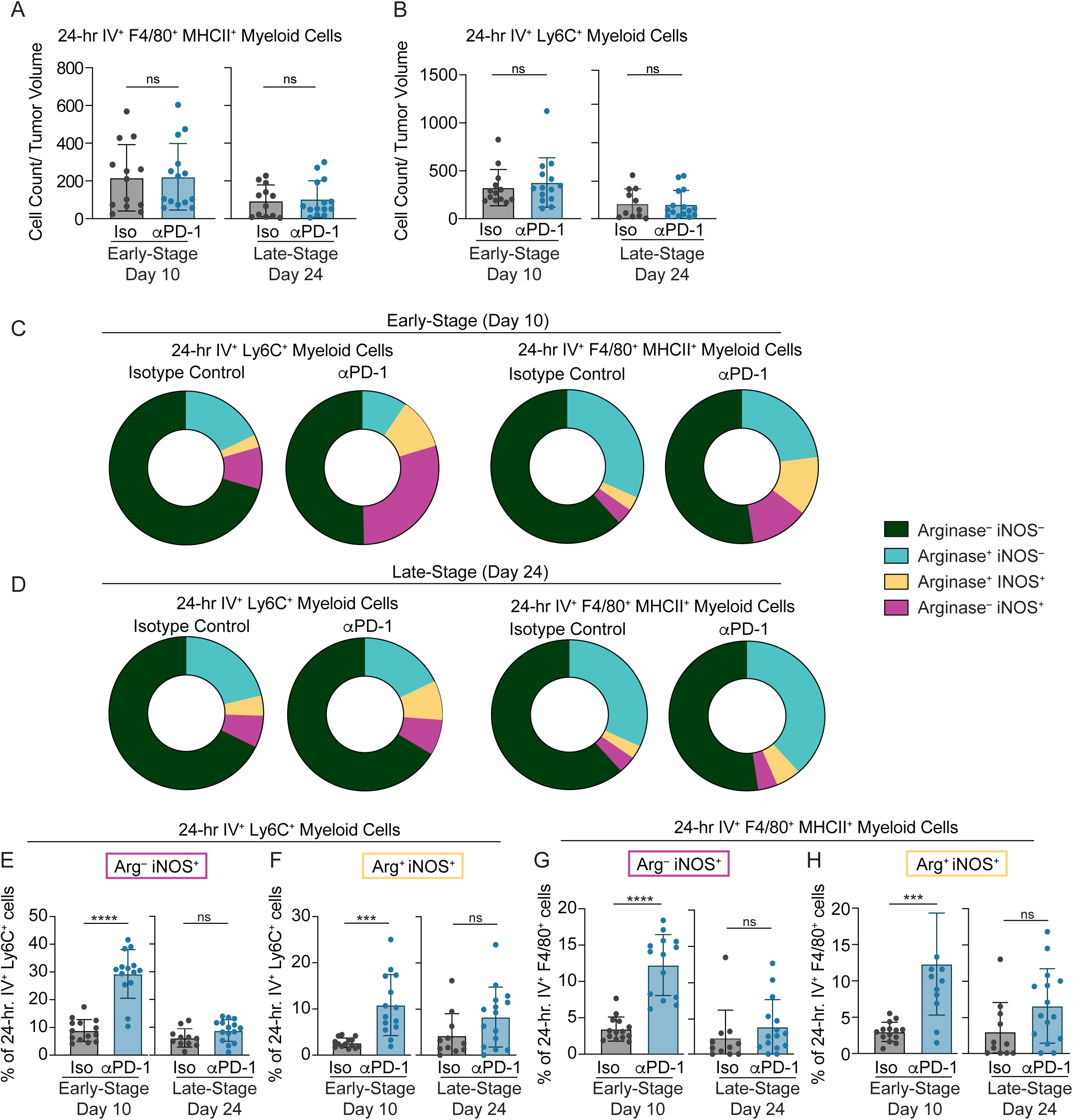
PD-1 blockade is more effective at inducing iNOS expression in myeloid populations in tumors at earlier stages of disease than later stages. Mice setup using the same experimental schematic as **Fig. 3A**. Female C57BL/6 mice were injected subcutaneously with 500,000 MC38 cells and randomized into isotype or anti-PD-1 treatment cohorts. ICI given on day 7 for the early-stage (followed by harvest at day 10) or day 18 and 21 for the late-stage (followed by harvest at day 24). 24 hours and 3 minutes prior to harvest, mice were injected with anti-CD45.2 antibodies conjugated to different fluorophores IV. F4/80^+^ MHCII^+^ myeloid cells are gated as F4/80^+^, Ly6G^−^, Ly6C^−^, MHCII^+^, CD11b^+^, while Ly6C^+^ myeloid cells are gated as F4/80^+^, Ly6G^−^, Ly6C^+^. A. Total cell count of recently recruited (24-hr IV^+^ 3-min IV^−^) F4/80^+^ MHCII^+^ myeloid cells normalized to tumor volume at the indicated harvest time points. B. Total cell count of recently recruited (24-hr IV^+^ 3-min IV^−^) Ly6C^+^ myeloid cells normalized to tumor volume at the indicated harvest time points. Data shown in panels **Fig. 4A**–**4B** are a combination of two independent experiments with 7-8 mice per experiment. C-D. Breakdown of the composition of recently recruited (24-hr IV^+^ 3-min IV^−^) Ly6C^+^ myeloid and F4/80^+^ MHCII^+^ myeloid cells that express arginase and/or iNOS at the indicated harvest time points. Comparison of the frequency of recently recruited (24-hr IV^+^ 3-min IV^−^) Ly6C^+^ myeloid cells expressing iNOS alone (E) or iNOS and arginase (F). Comparison of the frequency of recently recruited (24-hr IV^+^ 3-min IV^−^) F4/80^+^ MHCII^+^ myeloid cells expressing iNOS alone (G) or iNOS and arginase (H). Data shown in all panels are a combination of two independent experiments with 7-8 mice treatment group per experiment and are representative of 2-4 independent experiments. Statistical analyses in **Fig. 4A-4B** and **4E**–**4H** performed using unpaired Student’s t tests. In the Figure, ns indicates p>0.05, * indicates p<0.05, ** indicates p<0.01, and *** indicates p<0.001.

### Reducing CD8^+^ T cell trafficking to tumors impairs the protective effects of anti-PD-1 on tumor growth and reduces myeloid cell induction of iNOS

We next examined how disrupting lymphocyte trafficking to tumors impaired the protective effects of PD-1 inhibitors, leukocyte entry rate, and myeloid cell remodeling in the TME. To do this, we utilized the pharmacological agent FTY720, which targets the sphingosine-1-phosphate (S1P)/sphingosine-1-phosphate receptor 1 (S1PR1) axis required for lymphocyte egress from lymph nodes. Previous work has shown that FTY720 can trap T cells in lymph nodes by preventing S1P/S1PR1-mediated egress, causing a relative loss of T cells in the blood and the subsequent inability to traffic to tissues [42, 43]. Moreover, prior studies have used this approach to show that lymphocyte migration from the periphery to tumors is important for the therapeutic efficacy of PD-1 blockade [44–47]. Building on this prior work, we determined how FTY720 impacted leukocyte entry rates into the tumor as well as the phenotype of leukocytes in the TME. Mice were given MC38 tumor cells s.c. in the flank and FTY720 was administered intraperitoneally daily beginning at day 6, while anti-PD-1 or its isotype control were delivered intraperitoneally at days 8 and 11 (**Fig. 5A**). Mice received anti-CD45.2 IV at 24 hours and 3 minutes prior to harvest to label recently recruited leukocytes (24-hr IV^+^ 3-min IV^−^) (**Fig. 5A**). As expected, the administration of FTY720 abolished the protective effects of PD-1 inhibitors, with the tumor volumes of anti-PD-1 plus FTY720 being significantly higher than anti-PD-1 treated without FTY720 and not significantly different from the isotype-control treated animals (**Fig. 5B**).

**Figure 5:**
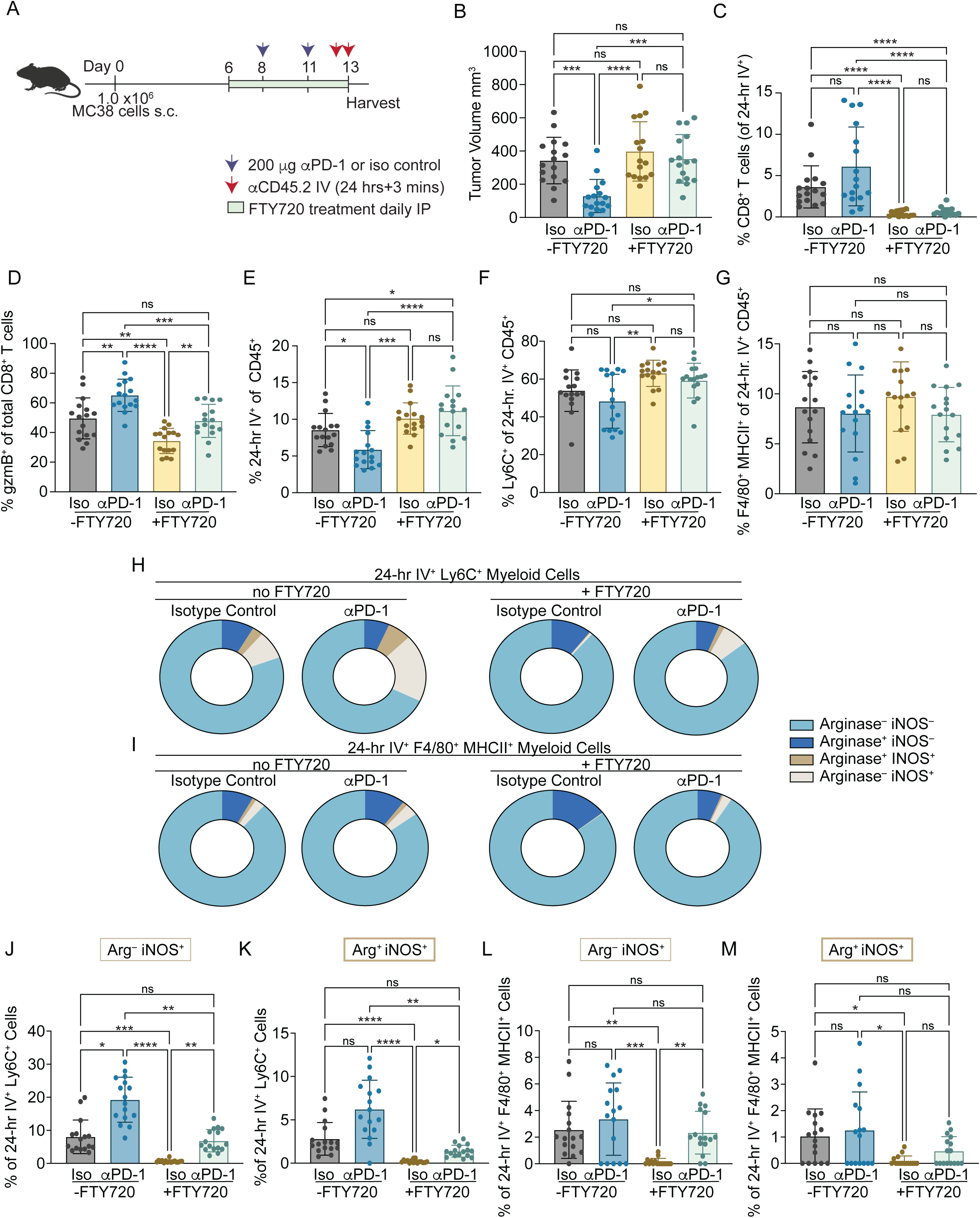
Modulating CD8^+^ T cell trafficking to tumors impairs the protective effects of anti-PD-1 on tumor growth, reduces CD8^+^ T cell recruitment, and reduces myeloid cell induction of iNOS. **A**. Experimental design to test how FTY720 administration impacts PD-1 ICI response. WT female C57BL/6 mice were injected subcutaneously with 750,000-1,000,000 MC38 cells and randomized into treatment cohorts. FTY720 was given via intraperitoneal injection daily from days 6-12 and ICI was given via intraperitoneal injection on days 8 and 11. 24 hours and 3 minutes prior to harvest (harvest on day 13), mice were injected intravenously with anti-CD45.2 antibodies conjugated to different fluorophores IV. B. Tumor volume at time of harvest. C. Percentage of CD8^+^ T cells within the 24-hr IV^+^ 3-min IV^−^ CD45^+^ population between treatment groups. Cells are gated on live, CD45^+^, Thy1.2^+^, CD8^+^. D. Percentage of Granzyme B^+^ CD8^+^ T cells. Cells are gated on live, CD45^+^, Thy1.2^+^, CD8^+^ (not further subgated on IV antibody). E. Percentage of 24-hr IV^+^ 3-min IV^−^ cells of the total CD45^+^ cells. F – M characterize myeloid subsets, where F4/80^+^ myeloid cells are gated as F4/80^+^, Ly6G^−^, Ly6C^−^, MHCII^+^, and CD11b^+^, and Ly6C^+^ myeloid cells are gated as: F4/80^+^, Ly6G^−^, Ly6C^+^. F. Percentage of Ly6C^+^ myeloid cells within the 24-hr IV^+^ 3-min IV^−^ CD45^+^ population. G. Percentage of F4/80^+^ MHCII^+^ myeloid cells within the 24-hr IV^+^ 3-min IV^−^ CD45^+^ population. H-I. Breakdown of the composition of 24-hr IV^+^ 3-min IV^−^ F4/80^+^ MHCII^+^ myeloid cells and Ly6C^+^ myeloid cells that express arginase and/or iNOS between treatment groups. Frequency of 24-hr IV^+^ 3-min IV^−^ Ly6C^+^ myeloid cells that express iNOS alone (J) or iNOS and arginase (K). Frequency of 24-hr IV^+^ 3-min IV^−^ F4/80^+^ MHCII^+^ myeloid cells expressing iNOS alone (L) or iNOS and arginase (M). Data shown in all panels are combined from two independent experiments with 8 mice per treatment group per experiment. Statistical analyses in **Fig. 5B-5G** and **5J-5M** performed using One-way ANOVA with multiple comparisons tests. In the Figure, ns indicates p>0.05, * indicates p<0.05, ** indicates p<0.01, *** indicates p<0.001, and **** indicates p<0.0001.

We next assessed how FTY720 impacted anti-tumor immunity with or without anti-PD-1 in the tumor-draining lymph node (TDLN), blood, and tumor. We predicted that PD-1 inhibitors would induce the expansion of tumor-specific CD8^+^ T cells in the TDLN and blood, consistent with previous work [44–46, 48], and that FTY720 would prevent these T cells from leaving the TDLN and subsequently entering the tumor. FTY720 caused a reduction in the overall abundance of T cells in the blood (**Fig. S5A, S5B**), as expected based on previous reports [42, 43]. We next assessed the impact of FTY720 on tumor-reactive CD8^+^ T cell responses in the periphery with or without anti-PD-1 by utilizing peptide/MHC tetramer reagents to identify tumor-antigen specific CD8^+^ T cells. Tetramer reagents are important for identify tumor-antigen specific CD8^+^ T cells in the periphery, where the overwhelming majority of T cells are not tumor-reactive [49]. In this study, we used tetramer reagents containing the Rpl18 neoantigen that has been described for MC38 (the peptide sequence KILTFDRL bound to the H2-K^b^ MHC class I molecule)[50]. Treatment with PD-1 ICI caused a trend towards an increase in the frequency of Rpl18-specific CD8^+^ T cells in the TDLN (**Fig. S5C, S5D**) and a significant increase in the frequency in the blood (**Fig. S5E**). Moreover, anti-PD-1 induced a significant upregulation of granzyme B (**Fig. S5F**) in the Rpl18-specific CD8^+^ T cell population in the TDLN, suggesting that PD-1 inhibitors are increasing the effector activity of tumor-specific CD8^+^ T cells in the periphery. In the TDLN, mice that received FTY720 showed higher levels of granzyme B in the anti-PD-1 treated group compared to isotype control-treated mice, similar to what was observed with anti-PD-1 treatment without FTY720 (though less granzyme B than mice treated with anti-PD-1 without FTY720, **Fig. S5F**). FTY720 caused a reduction in the frequency of Ki-67^+^ tumor-specific CD8^+^ T cells in both the isotype and anti-PD-1-treated groups (**Fig. S5G**), which may be reflective of trapping of proliferating T cells in the TDLN to a point where they have started exiting cell cycle. These data support a model where PD-1 inhibitors act on tumor-specific T cells in the TDLN, inducing numerical expansion, acquisition of effector features, and mobilization into the blood to traffic to tumors.

We next examined how FTY720 impacted recruitment to the tumor using the IV labeling approach. At the day 13 harvest time point, we observed that the frequency of CD8^+^ T cells within the 24-hr IV^+^ 3-min IV^−^ fraction was increased following PD-1 blockade compared to isotype control-treated mice (**Fig. 5C**), though this did not reach significance. FTY720 substantially reduced the abundance of CD8^+^ T cells within 24-hr IV^+^ 3-min IV^−^ fraction with or without PD-1 inhibitors (**Fig. 5C**), demonstrating the effectiveness of FTY720 to reduce T cell migration to tumors. Following FTY720 treatment, the overall frequency of granzyme B^+^ cells within the total CD8^+^ fraction within tumors was diminished (**Fig. 5D**), showing that the inability to recruit peripheral CD8^+^ T cells reduced the overall effector potential of the CD8^+^ T cell compartment in the tumor. Interestingly, however, FTY720 did not reduce the overall frequency of recently recruited leukocytes based on total CD45^+^ cells (**Fig. 5E**). We speculated that the lack of a difference between groups with and without FTY720 related to the inability of FTY720 to impact the trafficking of myeloid populations to the tumor, which comprise the majority of the recently recruited infiltrate in MC38 tumors both with and without anti-PD-1 (**Fig. S4A**). Indeed, FTY720 did not impact the frequency of Ly6C^+^ myeloid cells (**Fig. 5F**) or F4/80^+^ MHCII^+^ myeloid cells (**Fig. 5G**) within the 24-hr IV^+^ 3-min IV^−^ fraction in MC38 tumors. Importantly, despite not showing a reduction in myeloid cell entry, mice treated with FTY720 (both with and without PD-1 inhibitors) showed less induction of iNOS in the Ly6C^+^ and F4/80^+^ MHCII^+^ myeloid compartments (**Fig. 5H-5M**). The proinflammatory arginase^−^ iNOS^+^ compartment was virtually abolished with FTY720 in the isotype-treated mice and greatly diminished when comparing the effects of PD-1 blockade on proinflammatory cell recruitment with or without FTY720 (**Fig. 5J**, **5L**). We observed similar changes to the iNOS^+^ fractions within IV^−^ myeloid cells, indicating that a lack of T cell recruitment impacts myeloid cells in the local TME as well (**Fig. S5H-S5M**). These results support a model where lymphocyte recruitment to tumors is important for protective anti-tumor immunity following PD-1 inhibitors, and that both full induction of granzyme B in CD8^+^ T cells and remodeling of the myeloid compartment to a more proinflammatory phenotype is dependent on lymphocyte trafficking into tumors.

### PD-1 deletion in CD8^+^ T cells is sufficient to promote proinflammatory myeloid recruitment and create an immunostimulatory TME

The results from our FTY720 experiments suggested that PD-1 inhibitors acting on lymphocytes were playing a crucial role in the induction of proinflammatory programs in myeloid cells in the tumor. To test this hypothesis further, we determined whether genetic loss of PD-1 exclusively on CD8^+^ T cells was sufficient to drive the proinflammatory remodeling of the myeloid compartment. To do this, we utilized a CD8-specific PD-1 knockout (KO) mouse that has been previously described [51], which were generated by breeding mice containing a *loxP*-flanked *Pdcd1* gene [52] to mice expressing the gene encoding Cre recombinase under the control of the E8i enhancer element [53], referred to here as “CD8-PD-1^KO^ mice” (**Fig. 6A**). Previous work using CD8-PD-1^KO^ mice had shown that loss of PD-1 on CD8^+^ T cells was sufficient for protective immunity against MC38 [51]. Consistent with this previous work, we observed a trend towards reduced MC38 tumor size in CD8-PD-1^KO^ mice compared to littermate controls (PD-1^f/f^ mice lacking the E8i-cre transgene, referred to as wildtype (WT) mice here), though the results were not significantly different at this time point (**Fig. 6B**). Using this model, we asked how CD8^+^ T cell-restricted loss of PD-1 impacted leukocyte trafficking to tumors and functionality within the tumor. We observed an increase in the frequency of CD8^+^ T cells within the 24-hr IV^+^ fraction (**Fig. 6C**) and the density of recently recruited CD8^+^ T cells (**Fig. 6D**), supporting a cell-intrinsic role for PD-1 in regulating CD8^+^ T cell trafficking to tumors. In terms of the IV^−^ compartment, we observed an increase in the number of CD8^+^ T cells in the CD8-PD-1^KO^ mice compared to WT, but not the total number of CD45^+^ cells, F4/80^+^ MHCII^+^ myeloid cells, or Ly6C^+^ myeloid cells (**Fig. S6A-S6D**). As expected based on our FTY720 experiments, we did not observe a significant difference in the abundance of Ly6C^+^ or F4/80^+^ MHCII^+^ myeloid cells within the 24-hr IV^+^ 3-min IV^−^ compartment (**Fig. 6E**, **6F**). However, when PD-1 loss was restricted to CD8^+^ T cells, we observed an increase in iNOS induction in both the Ly6C^+^ or F4/80^+^ MHCII^+^ myeloid cells within the 24-hr IV^+^ 3-min IV^−^ compartment (**Fig. 6G**, **6H**) as well as the IV^−^ compartment (**Fig. S6E**). There was also a significant increase in the frequency of arginase^−^ iNOS^+^ cells within both recently recruited myeloid populations, and a slight but not significant increase in the frequency of arginase^+^ iNOS^+^ cells (**Fig. 6I-6K**). Within the IV^−^ myeloid cells, the CD8-PD-1^KO^ mice also had an elevated frequency of iNOS^+^ cells compared to the WT control (**Fig. S6F-S6I**), suggesting that changes downstream of PD-1 loss in CD8^+^ T cells drive remodeling of the myeloid compartment to be more proinflammatory in MC38 tumors. These data support a model where PD-1 loss exclusively on CD8^+^ T cells is capable of driving increased migration to tumors, and support a role for PD-1 modulation on CD8^+^ T cells being a driver of myeloid cell remodeling in tumors.

**Figure 6:**
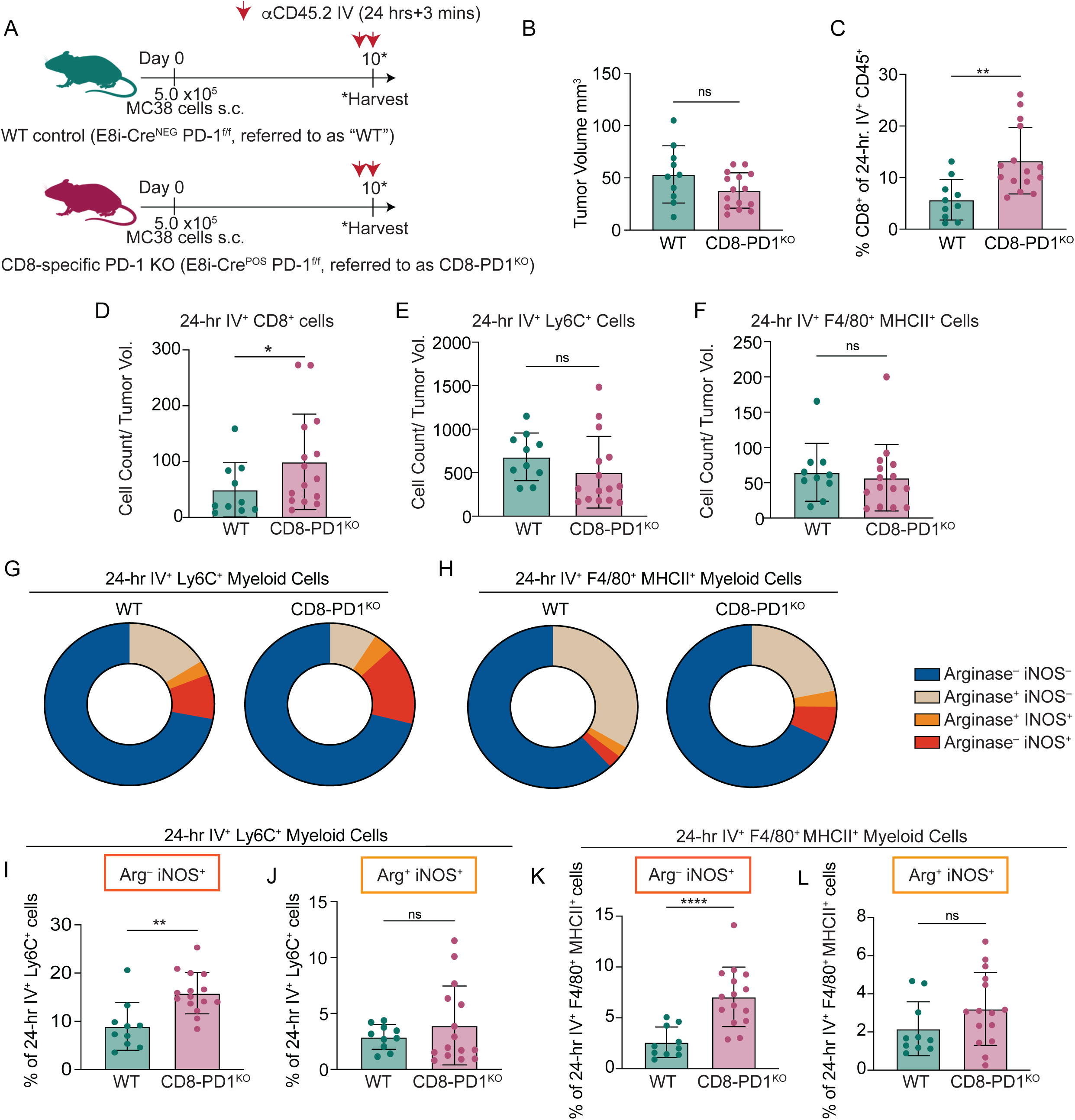
PD-1 deletion in CD8^+^ T cells is sufficient to increase inflammatory myeloid recruitment and create an immunostimulatory TME. **A.** Experimental design to test the role of CD8-specific PD-1 deletion on trafficking and myeloid remodeling in MC38 tumors. Previously described CD8-specific PD-1 KO mice [51] (E8i-Cre^POS^ PD-1^f/f^, referred to as CD8-PD-1^KO^) or littermate controls (E8i-Cre^NEG^ PD-1^f/f^, referred to as WT) were injected subcutaneously with 500,000-1,000,000 MC38 cells and harvested on day 10 post-tumor cell injection. 24 hours and 3 minutes prior to harvest, mice were injected with anti-CD45.2 antibodies conjugated to different fluorophores IV. B. Tumor volume at time of harvest. In C – F, cells are gated on live, CD45^+^. C. Percentage of CD8^+^ T cells within the live, CD45^+^, 24-hr IV^+^ 3-min IV^−^ population. CD8^+^ T cells were gated at Thy1.2^+^ CD8^+^. D. Total cell count of 24-hr IV^+^ 3-min IV^−^ CD8^+^ T cells normalized per unit tumor volume. For myeloid analyses in E – L, F4/80^+^ MHCII^+^ myeloid cells are gated as F4/80^+^, Ly6G^−^, Ly6C^−^,MHCII^+^, and CD11b^+^, while Ly6C^+^ myeloid cells gated as F4/80^+^, Ly6G^−^, Ly6C^+^. E. Total cell count of 24-hr IV^+^ 3-min IV^−^ Ly6C^+^ myeloid cells normalized per unit tumor volume. F. Total cell count of 24-hr IV^+^ 3-min IV^−^ F4/80^+^ MHCII^+^ myeloid cells normalized per unit tumor volume. G-H. Frequency of 24-hr IV^+^ 3-min IV^−^ F4/80^+^ MHCII^+^ myeloid cells and Ly6C^+^ myeloid cells that express arginase and/or iNOS between treatment groups. Frequency of 24-hr IV^+^ 3-min IV^−^ Ly6C^+^ myeloid cells expressing iNOS alone (I) or iNOS and arginase (J). Frequency of 24-hr IV^+^ 3-min IV^−^ F4/80^+^ MHCII^+^ myeloid cells expressing iNOS alone (K) or iNOS and arginase (L). Data shown in all panels are a combination of two independent experiments with 4-7 mice per treatment group per experiment. Statistical analyses in **Fig. 6B-6F** and **6I**–**6L** performed using unpaired Student’s t tests. In the Figure, ns indicates p>0.05, * indicates p<0.05, and ** indicates p<0.01.

## Discussion

To date, many studies have relied on perturbing migration to show the importance of leukocyte recruitment in driving productive anti-tumor immunity, including the use of the drug FTY720 [44–47]. While these studies have shown that migration from the periphery to tumors can be a key driver of protective anti-tumor immunity following PD-1 ICI, the inability to granularly distinguish recruited leukocytes from tumor-residing populations has been a major limitation for the field and has contributed to a significant gap in knowledge regarding the normal physiology of recently recruited populations in the unperturbed, native tissue microenvironment. Our study provides new insights into how PD-1 ICI impacts leukocyte recruitment to tumors, demonstrating that successful PD-1 ICI is associated with increased recruitment of CD8^+^ T cells to tumors which drives the remodeling of the intratumoral myeloid compartment to be more proinflammatory. Importantly, PD-1 ICI given at later stages of disease fails to induce these changes and fails to drive effective tumor control. This work provides insights into the phenotype of recently recruited leukocytes and the impact of recently recruited leukocytes on intratumoral leukocyte populations, providing further rationale to target recruitment to improve outcomes for patients.

The degree of CD8^+^ T cell infiltration into tumors in patients has consistently been shown to be associated with enhanced overall survival and response to ICI in multiple cancer types [54–59]. It is well established that CD8^+^ T cells are contact dependent killers, so having greater quantities of these T cells in the TME would increase the probability of the immune system successfully eliminating the tumor [10]. However, T cell exhaustion in the TME reduces the quality of the CD8^+^ T cell response, thereby worsening the effective T cell-to-tumor cell ratio. PD-1 ICIs do little to impact the epigenetic landscape associated with T cell exhaustion [24, 60–64], suggesting that exclusively targeting CD8^+^ T cells in the TME may have limited therapeutic impact. Fortunately, intratumoral CD8^+^ T cells are not the only tumor-specific CD8^+^ T cells in the body that can be leveraged for therapeutic gain. In cancer, the TDLN contains a reservoir of functional CD8^+^ T cells [32, 46, 48, 65, 66]. Consequently, approaches to expand and mobilize these T cell populations from the periphery to the tumor could be more effective at driving productive anti-tumor immunity than exclusively targeting T cells in the TME. Indeed, recent work has demonstrated that peripheral immune cell activation is predictive of productive anti-tumor immunity after PD-1 inhibition [44, 45, 48, 67–70]. Moreover, studies in patients have shown that infiltration of tumors by new T cell clones from the blood after PD-1 ICI is associated with better response to therapy [71–74]. Our results show that at early stages of MC38, when PD-1 ICI is effective, PD-1 ICIs are capable of increasing migration of CD8^+^ T cells from the periphery to the tumor, and recently recruited CD8^+^ T cells displayed a less exhausted phenotype than their tumor-residing (IV^−^) counterparts. However, our results also show that PD-1 ICI is more effective at driving the recruitment of CD8^+^ T cells to tumors at earlier stages of disease than later stages of disease. We predict that the reduced ability to mobilize tumor-specific CD8^+^ T cells from the periphery to the tumor is a major mechanism contributing to resistance to ICI, and that developing combination therapies to better expand and mobilize CD8^+^ T cells from the periphery to the tumor will be important for overcoming this barrier to improve outcomes in patients.

Though CD8^+^ T cells play an important role in anti-tumor immunity following PD-1 ICI or deletion [30, 51, 75–77], CD8^+^ T cells are only one component of the immune infiltrate in tumors. Indeed, our results showed that the minority of the cells in the 24-hr IV^+^ 3-min IV^−^ infiltrate were CD8^+^ T cells. F4/80^+^ MHCII^+^ myeloid cells (likely macrophages) and Ly6C^+^ myeloid cells (likely monocytes) far outnumbered the CD8^+^ T cell population in the 24-hr IV^+^ 3-min IV^−^ infiltrate. Prior work has demonstrated the complex and dualistic nature of tumor-infiltrating myeloid cells. Indeed, sometimes myeloid cells can exert critical proinflammatory or anti-tumor functions, while other times these myeloid cells can be hijacked by tumors and suppress host immunity and/or promote tumor growth [7, 38, 39]. Therapeutically, myeloid cells represent a significant opportunity, as these cells are sensitive to changes within the TME and massive numbers can be recruited into the tumor from the peripheral blood. Indeed, PD-1 ICI has been shown to drive a remodeling of the macrophage compartment in responsive tumors, shifting these myeloid cells from a more immunosuppressive M2-like CX3CR1^+^ state into a more proinflammatory M1-like iNOS^+^ state [78, 79]. Our findings are consistent with these results, showing that at early stages of disease when ICI was more effective, anti-PD-1 drove an increase in iNOS within both the Ly6C^+^ and F4/80^+^ MHCII^+^ myeloid cell fractions. This increase in iNOS expression was noted in both the recently recruited (24-hr IV^+^ 3-min IV^−^) and IV^−^ fractions, suggesting a remodeling of the Ly6C^+^ and F4/80^+^ MHCII^+^ myeloid populations that were both trafficking into the tumor as well as already residing within the TME. Prior work has also demonstrated the importance of monocyte-T cell interactions within tumors, and that these interactions can facilitate potent anti-tumor immunity [13]. Because we observed concurrent T cell and Ly6C^+^ myeloid cell recruitment into tumors after ICI, we speculate that recruiting both populations may drive synergy in anti-tumor immunity, consistent with previous work [13]. Consistent with this hypothesis, when FTY720 was given, we observed reduced iNOS induction in the myeloid compartment, suggesting that CD8^+^ T cell recruitment drives myeloid cell remodeling towards a more proinflammatory phenotype. Our data show also support a model where some immunostimulatory myeloid cells may be maturing locally. The local maturation of IV^−^ myeloid cells may be driven by changes in the local inflammatory milieu, as previous work has shown that Type I [80] or Type II IFN [78] can lead to the induction of the proinflammatory phenotype in myeloid cells following ICI in tumors. Considering the plasticity of myeloid cells, future studies examining how different factors in the TME shape myeloid cell differentiation and function upon tumor entry are warranted.

To date, the field has been severely lacking in tools to timestamp leukocytes upon entering tissues, limiting our understanding of leukocyte entry rates over time and the cellular and molecular underpinnings of tissue adaptation and function. We contend that the IV antibody labeling approach provides an alternative to the photoactivatable system [13] currently used for these types of studies in that extensive breeding is not needed, multiple time points can be labeled within a single mouse, and it is not limited by issues of tissue penetration [14, 15]. Our results obtained using the IV labeling method are consistent with work performed in MC38 tumors using the photoactivatable system, including the observation that T cell entry into MC38 tumors dynamically persisted over time and PD-1 ICI impacted both recently recruited and non-recently recruited T cell populations [33]. Despite the advantages of the IV labeling approach, there are limitations that should be considered when implementing this method. The IV labeling approach only labels the leukocytes in the blood for a narrow window of time and does not counter-label cells that were in the tumor prior to labeling. We speculate that the fraction of leukocytes that could have entered the blood after the 24-hour labeling window and then entered the tumor (and therefore would be in the IV^−^ fraction) is small, but this is a current limitation that should be considered when interpreting the data. Additionally, the fluorescent antibody-based approach provides a relatively narrow window to identify recently recruited cells [14, 15]. Future work combining the IV labeling approach with more durable labeling methods like fate mapping [81] could be useful for longer-term studies.

Considering the importance of migration for both innate and adaptive leukocyte subsets, we contend that leukocyte migration is a critical axis to consider when designing new immunotherapy approaches to improve outcomes in patients. We predict that patients who respond well to ICI are capable of mobilizing enough of the right immune cells to the tumor to eliminate malignant cells, while patients who do not respond well to ICI could benefit from combination approaches aiming to expand and mobilize the right immune cells to their tumors. We believe that the IV antibody labeling method could provide an efficient way to screen drugs for their ability to promote the migration of key leukocyte subsets to tumors in the preclinical setting, providing a rationale for how to improve leukocyte migration in patients. Ultimately, expanding the use of existing modalities [82, 83] and developing novel agents to label circulating leukocytes prior to tissue entry in patients could be an effective way to use this approach in translational studies to ask these questions directly in patients.

## Materials and Methods

### Mice

Wildtype female C57BL/6 (JAX stock number 000664) mice were purchased from The Jackson Laboratory. The CD8-specific PD-1^f/f^ mice were donated to the Pauken laboratory by Dr. Arlene H. Sharpe (Harvard Medical School) and have been previously described [51]. Briefly, mice purchased from the Jackson Laboratory containing a Cre-IRES-GFP cassette under the control of the CD8a E8i enhancer (full name C57BL/6-Tg(Cd8a-cre)1Itan/J, JAX stock number 008766) [53] were bred to mice containing a *loxP*-flanked *Pdcd1* allele (termed PD-1^f/f^ mice) that had been previously described [52]. In this study, mice were bred with one copy of cre and two copies of the *loxP*-flanked *Pdcd1* gene (E8i-cre^POS^ PD-1^f/f^, referred to as CD8-PD-1^KO^) to no copies of cre and two copies of the loxP-flanked *Pdcd1* gene so the littermates could be used as PD-1-sufficient controls (E8i-cre^NEG^ PD-1^f/f^ referred to as WT controls), consistent with previous work [51]. Genotyping was performed using PCR on tail biopsies collected at 21 days of age or earlier using Transnetyx Genotyping Services (Transnetyx). Tumor cells were implanted into female mice between 7-12 weeks of age. All mice were maintained in specific pathogen-free facilities at The University of Texas MD Anderson Cancer Center (MD Anderson) under standard housing, husbandry, and diet conditions in accordance with Institutional Animal Care and Use Committee (IACUC) and National Institutes of Health (NIH) guidelines. Animal protocols were approved by the IACUC at MD Anderson (under the Animal Care and Use Form numbers 00002324-RN00 and 00002324-RN01) prior to performing the work.

### Tumor cell lines, implantation, and monitoring

Tumor cell lines were cultured in a 37°C incubator with 5% CO_2_, and cells were harvested at passage 2-3 after thaw. To remove tumor cells from tissue culture flasks, the tumor cells were first washed with pre-warmed phosphate buffered saline (PBS, without calcium or magnesium, Corning), incubated briefly in 0.05% trypsin-ethylenediaminetetraacetic acid (EDTA) (Gibco) to remove from the flask, and washed with pre-warmed media (the media used is specific for each cell line, as described). MC38 colon adenocarcinoma cells (donated by Dr. Arlene H. Sharpe, Harvard Medical School) were cultured in Dulbecco’s Modified Eagle Medium (DMEM) (cat. 11995-065 from Gibco, which contains 4.5 g/L D-glucose, L-glutamine, and 110 mg/L sodium pyruvate) supplemented with 10% fetal bovine serum (FBS; Gibco) and 100 U penicillin and 100 μg streptomycin (Gibco). The B16-OVA melanoma cell line was purchased from Millipore Sigma. B16-OVA was cultured in the same medium as MC38, but after the first passage was also supplemented with G418 (Gemini Bio) at a concentration of 1 mg/ml to select for cells expressing OVA per the manufacturer’s instructions. The YUMM3.3 cell line was purchased from ATCC. YUMM3.3 tumor cells were cultured in DMEM-F12 medium (Gibco) supplemented with 10% FBS, 100 U penicillin, 100 μg streptomycin, 1X MEM non-essential amino acids (Corning), 1X Gluta-MAX (Gibco), 0.01 M HEPES (also known as 4-(2-hydroxyethyl)-1-piperazineethanesulfonic acid; Gibco), and 1 mM sodium pyruvate (Gibco). A total of 2.5×10^5^ - 1×10^6^ cells were injected subcutaneously into the flank skin on the shaved abdomen of mice anesthetized with inhaled isoflurane.

Tumors were measured every 2-3 days with calipers beginning at day 7-8. Tumor volume was calculated using the formula ½ x D x d^2^, where D is the major axis and d is the minor axis. The time point of tumor harvest is indicated in the Figure Legends. Due to the time points analyzed, mice typically reached experimental endpoints prior to reaching primary or humane endpoints. However, mice were closely monitored and were euthanized if primary or humane endpoints were reached in accordance with IACUC guidelines and our approved IACUC protocols. Primary endpoint was a tumor volume of 2000 mm^3^. Humane endpoints included: 1) tumors ulcerated to 2 mm or greater or becoming necrotic, infected, or displaying signs of trauma, 2) touching of the tumor induced pain (which was evaluated using vocalization, flinching, or withdrawal from the mouse), 3) the tumor interfered with normal daily activities including the ability to eat or drink, or 4) a severely deteriorating body condition (including 20% weight loss, anorexia, rapid or slow breathing, dermatitis, diarrhea, urinary incontinence, hunched posture, ruffled fur, lethargy, and self-induced trauma).

### Therapeutic agent administration

Timing for therapeutic agents given are indicated in the relevant Figures and Figure Legends. Mice received 200 μg/dose (in 200 μl of sterile PBS) of anti-PD-1 antibodies (clone 29F.1A12, BioXCell or Leinco) or its isotype control (Rat IgG2a clone 2A3, BioXCell) intraperitoneally. For early-stage experiments, mice received a single dose of therapeutic antibodies at day 7. For late-stage experiments, mice received two doses of therapeutic antibodies, one at day 18 and one at day 21. For FTY720 (Millipore-Sigma) experiments, two doses of anti-PD-1 or isotype control were given, one at day 8 and one at day 11. Mice received FTY720 (25 μg/dose given intraperitoneally in 100 μl of sterile PBS) beginning at day 6 post-tumor implantation and continued daily until harvest.

### *In vivo* antibody labeling to time stamp leukocytes prior to entry into the tumor

*In vivo* antibody labeling was performed as described [14, 15]. Briefly, mice were anesthetized with inhaled isoflurane and given a retro-orbital injection of fluorescently conjugated anti-CD45.2 (clone 104) antibodies intravenously. For the 24-hr time point, 5 μg of antibody was administered in 100 μl of sterile PBS. For the 3-min time point, 2 μg of antibody was administered in 100 μl of sterile PBS. Antibodies were conjugated to AF647, AF488, BV421, or BV510 (Biolegend).

### Lymphocyte isolation from mouse tissues

TDLN were mechanically disaggregated through a 70 μm cell strainer. Spleens were mechanically disaggregated through a 70 μm cell strainer and then red blood cells were lysed using ACK Lysis Buffer (Gibco). Peripheral blood mononuclear cells (PBMCs) were isolated from peripheral blood using histopaque-1083 (Millipore-Sigma) gradients. Briefly, blood was collected into room temperature 4% sodium citrate (Millipore-Sigma, generated by diluting sodium citrate dihydrate into distilled water) as an anti-coagulant. After 5-10 minutes, Roswell Park Memorial Institute (RPMI) supplemented with 5% FBS was used to quench this reaction. Room temperature histopaque-1083 was underlaid, and white blood cells were separated from red blood cells via centrifugation.

MC38 and B16-OVA tumors were mechanically disaggregated using a GentleMACS (Miltenyi). YUMM3.3 tumors were disaggregated using fine chopping. The disaggregated tissue was digested with Collagenase Type I (400 U/ml; Worthington Biochemical) for 30 minutes at 37°C, followed by an additional dissociation step with the GentleMACS. Samples were washed using RPMI supplemented with 5% FBS and filtered through a 70 μm cell strainer. The single cell suspensions were immediately stained for flow cytometry.

### Flow Cytometry

Single cell suspensions were generated as described above and labeled with LIVE/DEAD Zombie NIR Dye (Biolegend) in PBS. Cells were then pre-incubated with TruStain Fc Receptor Block (anti-mouse CD16/CD32, clone 93, BioLegend), then labeled with extracellular antibodies in PBS supplemented with 2% FBS including: Thy1.2 (clone 30-H12), CD8α (clone 53-6.7); CD45 (clone 30-F11), PD-1 (clone RMPI-30), TIM-3 (clone RMT3-23), TIGIT (clone A17200C), CD160 (clone 7H1), LAG-3 (clone C9B7W), CD44 (clone IM7), MHCII (clone M5/114.15.2), CD11c (clone N418), CD11b (clone M1/70), Ly6G (clone 1A8), Ly6C (clone HK1.4), all from BioLegend, and/or F4/80 (clone T45-2342, from BD Biosciences). The H2-K^b^:Rpl18 tetramer (containing the peptide sequence KILTFDRL) was obtained from the NIH Tetramer Core Facility, and stained along with the extracellular antibodies. For analysis of intracellular markers, cells were permeabilized following the extracellular stain with the Foxp3/Transcription Factor Staining kit (ebioscience/ThermoFisher Scientific). Cells were subsequently stained with antibodies to detect the intracellular markers, including Granzyme B (clone GB11, BD Biosciences), Ki-67 (clone B56, BD Biosciences), TCF-1 (clone E6O1K, Cell Signaling), iNOS (clone A982, Miltenyi Biotec), Arginase (clone A1exF5, Biolegend), and/or CD206 (clone C068C2, Biolegend). Samples were acquired on either a X20 Fortessa (BD Biosciences) or Novocyte Opteon spectral cytometer (Agilent Technologies) and analyzed with FlowJo software (TreeStar). Inhibitory receptor co-expression analyses were performed in FlowJo software using the Boolean gating tool. PD-1, TIM-3, LAG-3, CD160, and TIGIT were included in the analysis for steady state mice, and TIM-3, LAG-3, CD160, and TIGIT were included in the analysis for isotype control-vs. anti-PD-1 treated mice. PD-1 was excluded from that analysis due to the potential for the in vivo anti-PD-1 antibody interfering with the flow cytometry antibody.

### Statistical Analysis

Statistics were performed with Prism software (GraphPad), and p-values of less than 0.05 were considered statistically significant. Prior to selecting the specific test to be used, normality was assessed using the D’Agostino-Pearson omnibus normality tests and Shapiro-Wilk normality tests. In cases where all groups were normally distributed, then statistical tests were selected to account for assuming Gaussian distribution. In the case where any group was not normally distributed, then statistical tests were selected to account for not assuming Gaussian distribution. For analyses involving only two groups, t-tests were performed. For unpaired observations, if data were normally distributed then an unpaired t test was performed, and if data were not normally distributed then a Mann-Whitney test was performed. For paired observations, if data were normally distributed then a paired t-test was performed, and if the data were not normally distributed then a Wilcoxon matched-pairs signed rank test was performed. If analyses contained more than two groups but only one independent variable, then a one-way ANOVA with post-test comparisons was typically performed. If data were normally distributed, then an ordinary one-way ANOVA was performed. If data were not normally distributed, then a Kruskal-Wallis test was performed. If analyses contained more than two groups and two independent variables, a two-way ANOVA with post-test comparison was performed. For multiple comparison tests, the mean rank of each column was compared with the mean rank of every other column. For each individual plot, the statistical test used is included in the Figure Legends. Asterisks indicating significance in the Figure Legends correspond to p<0.05 (*), p<0.01 (**), p<0.001 (***), and p<0.0001 (****).

## Conflicts of Interest Statement

KEP reports a patent on T cell exhaustion state-specific gene expression regulators and uses thereof (US2021/0309965 A1) that is unrelated to the current work. MMG reports a personal honorarium from Springer Nature Ltd (as a deputy editor for *Nature Precision Oncology*) and served as a paid consultant for Merck. The remaining authors indicate no other conflicts of interest to disclose. No payments or services were received from a third party in the last 36 months with entities that could be perceived to influence the submitted work.

## Acknowledgements

We would like to thank all the members of the Pauken laboratory and Schenkel laboratory for useful feedback on the project. We thank Dr. Arlene H. Sharpe (Harvard Medical School) for sharing the CD8-specific PD-1 KO mouse. This work was supported in part by institutional start-up funds (The University of Texas MD Anderson Cancer Center) to K.E.P., the Melanoma SPORE Developmental Research Program at The University of Texas MD Anderson Cancer Center (supported by P50CA221703 from the National Cancer Institute (NCI) of the NIH) to K.E.P., and the R01 award R01CA282027 from NCI to M.M.G. K.E.P. and J.M.S. were Andrew Sabin Family Foundation Fellows at MD Anderson and were supported by a University of Texas Rising STARs Award. J.M.S. was supported by the NCI grant K08-CA256044 and the Physician Scientist Program at MD Anderson Cancer Center. J.M.S. is a Cancer Prevention and Research Institute of Texas (CPRIT) Scholar in Cancer Research and was supported by the University of Texas Rising STARs Award. M.M.G. was a CPRIT Scholar in Cancer Research. This research is supported in part by The University of Texas MD Anderson Cancer Center Support Grant CA016672 including animal housing and care in the Research Animal Support Facility. We thank the NIH Tetramer Core Facility (NIH Contract 75N93020D00005 and RRID:SCR_026557) for providing the H2-K^b^:Rpl18 tetramer (containing the peptide sequence KILTFDRL) which was used in this study.

## Author Contributions

OA and KEP conceptualized the study. OA, JMS, and KEP contributed to the methodology development. OA led the investigation, data generation, data curation, visualization, data analysis, and data interpretation. AJS, AMS, and AKS also contributed to investigation and data generation. SSW, MMG, JMS, and KEP contributed to data interpretation. OA and KEP wrote the original draft of the manuscript. JMS, MMG, SSW, AMS, and AJS revised the manuscript. All authors approved the manuscript. KEP led the funding acquisition, supervision, and project administration for the manuscript.

---

**Figure S1.**
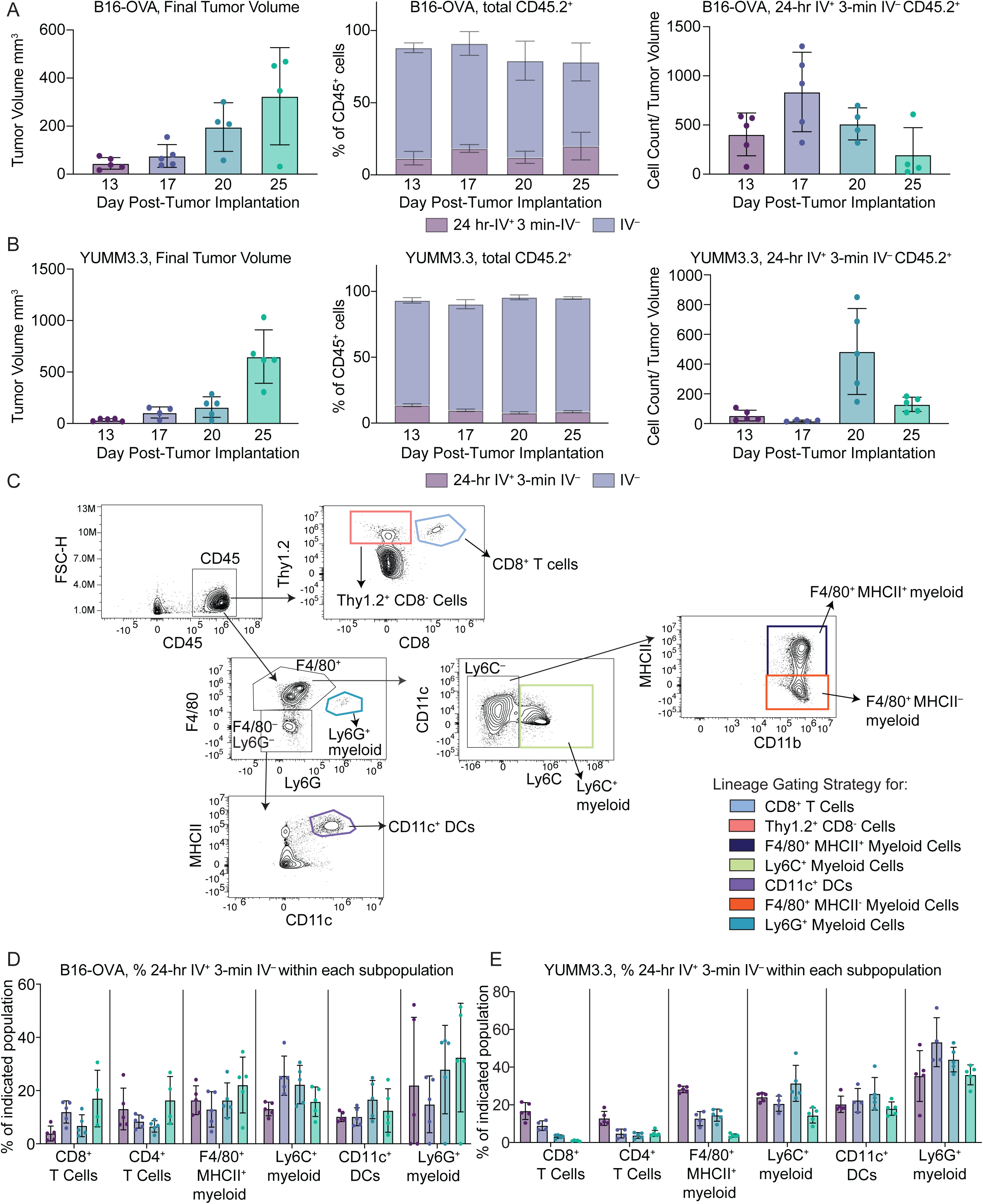
IV antibody labeling in B16-OVA and YUMM3.3 melanoma lines and gating strategy for immune cell lineages. IV antibody labeling in B16-OVA (A) vs. YUMM3.3 melanoma (**B**) injected into WT C57BL/6 mice. Mice received 250,000-500,000 tumor cells subcutaneously in the flank. In both **A** and **B**, left panel shows tumor volume at time of tumor harvest, middle panel shows the percentage of total CD45^+^ cells that were 24-hr IV^+^ 3-min IV^−^ vs. total CD45^+^ cells that were IV^−^, and the right panel shows the total cell count of 24-hr IV^+^ 3-min IV^−^ CD45^+^ immune cells normalized to tumor volume. **C.** Gating strategy for the major lineages shown throughout the manuscript. **D – E**. Quantification of the frequency of each indicated immune cell lineage that was recently recruited (24-hr IV^+^ 3-min IV^−^) over time in B16-OVA (**D**) and YUMM3.3 (**E**). Cells were gated down to each lineage first, and then the frequency of 24-hr IV^+^ 3-min IV^−^ was determined within each population. All lineages are gated on live, CD45^+^. Then, individual lineages further defined as: CD8^+^ T cells: Thy1.2^+^ CD8^+^; F4/80^+^ MHCII^+^ myeloid cells: F4/80^+^, Ly6G^−^, Ly6C^−^, MHCII^+^, CD11b^+^; Ly6C^+^ myeloid cells: F4/80^+^, Ly6G^−^, Ly6C^+^; CD11c^+^ dendritic cells (DCs): F4/80^−^, Ly6G^−^, CD11c^+^, MHCII^+^; Ly6G^+^ myeloid cells: F4/80^−^, Ly6G^+^. **Fig. S1A, S1B, S1D**, and **S1E** are representative of 2 independent experiments with 4-5 mice per group per experiment.

**Figure S2.**
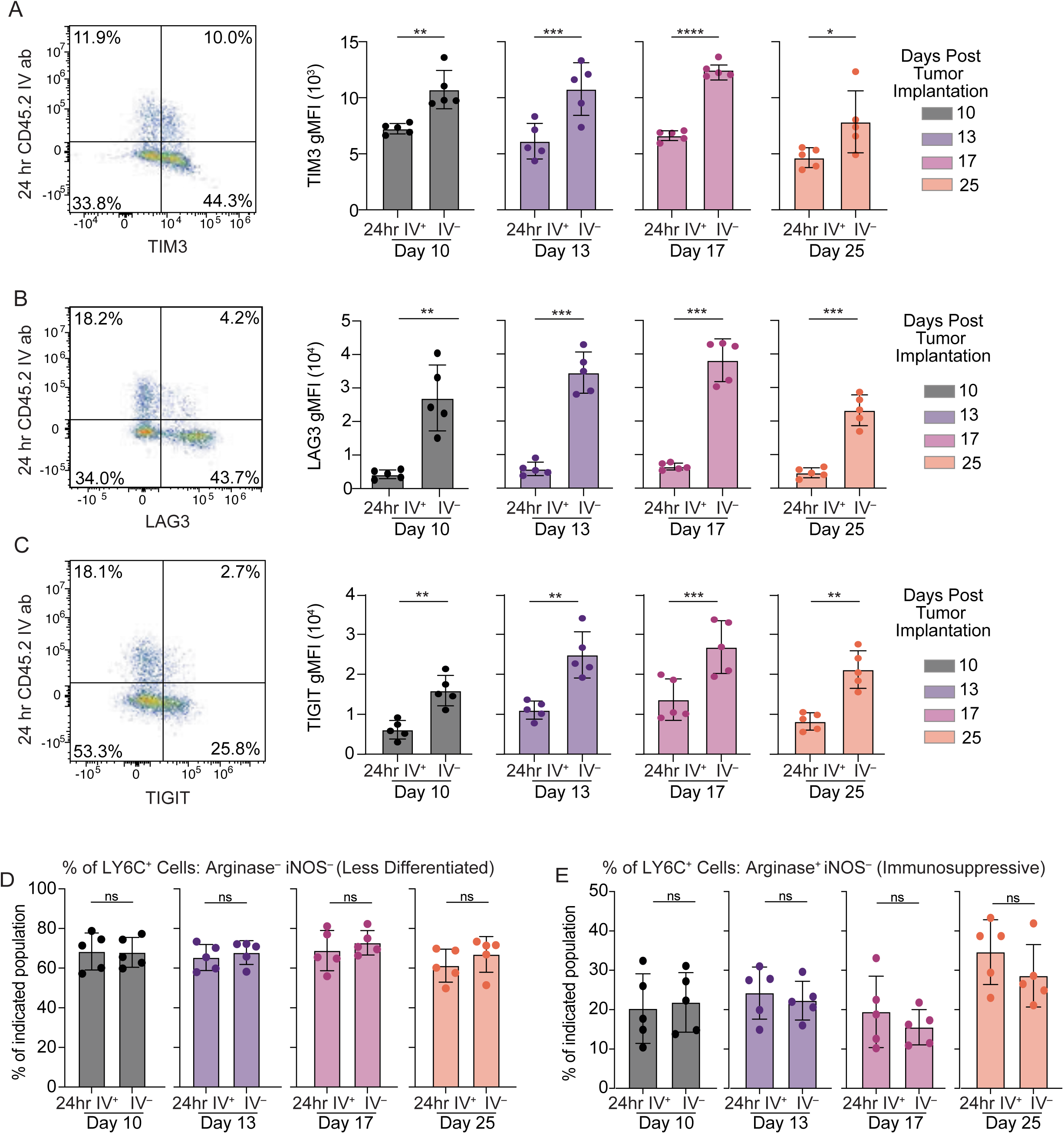
Inhibitory receptor expression on CD8^+^ T cells in MC38 tumors. Analyses shown are from the mice plotted in **Figure 2**. Plots are gated on live, CD45^+^, Thy1.2^+^, CD8^+^, 3-min IV^−^ to show comparisons of 24-hr IV^+^ vs. IV^−^ CD8^+^ T cells within the MC38 tumor tissue. “24-hr IV^+^” in the Figure therefore indicates “24-hr IV^+^ 3-min IV^−^”, while “IV^−^” indicates “24-hr IV^−^ 3-min IV^−^”. (Left) Representative flow plot showing indicated inhibitory receptors (TIM-3 (**A**), LAG-3 in (**B**), and TIGIT in (**C**)) vs. the 24-hr anti-CD45.2 label on CD8^+^ T cells in MC38 tumors (day 17). (Right) Quantification of each indicated inhibitory receptor expression using gMFI in the indicated CD8^+^ TIL populations. Data are from one experiment with n = 5 mice per group. The data are representative of two independent experiments with 3-5 mice per group per experiment. Quantification of the frequency of indicated populations of Ly6C^+^ myeloid cells that were (**D**) arginase^−^ and iNOS^−^ (e.g., less differentiated) or (**E**) arginase^+^ iNOS^−^ (more immunosuppressive) at different time points. Ly6C^+^ myeloid cells are gated as F4/80^+^, Ly6C^+^, Ly6G^−^. Data shown in **S2D and S2E** are from one experiment with n = 5 mice per group. The data are representative of two independent experiments with 4-5 mice per group per experiment. Statistics calculated using a paired Student’s t test at each individual time point. In the Figure, ns indicates p>0.05, * indicates p<0.05, ** indicates p<0.01, *** indicates p<0.001, and **** indicates p<0.0001.

**Figure S3.**
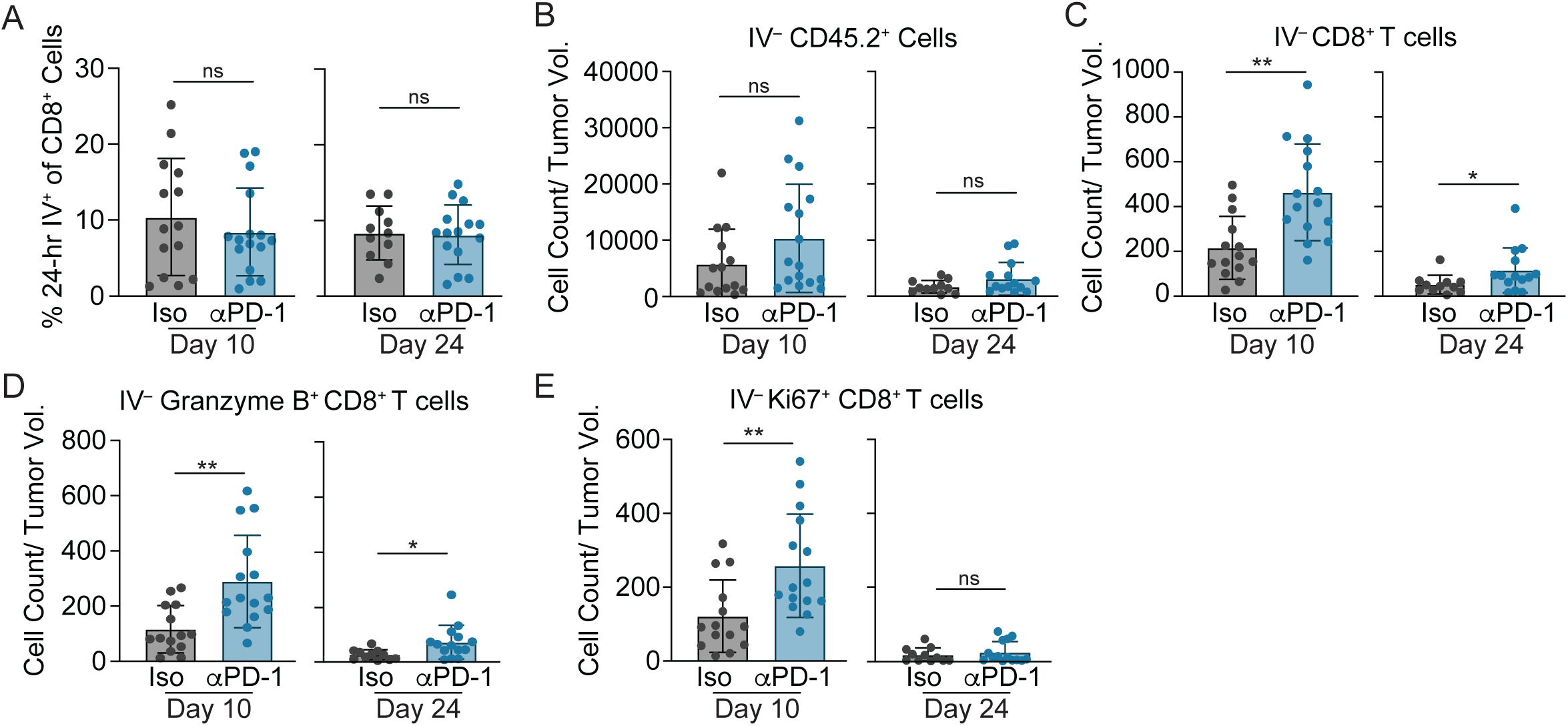
Impact of PD-1 ICI on CD8^+^ T cells in the IV^−^ compartment following early vs. late treatment in MC38 tumors. **A.** Frequency of the CD8^+^ T cell population (gated on live, CD45^+^, Thy1.2^+^, CD8^+^) that is 24-hr IV^+^ 3-min IV^−^ (gated on lineage first to determine the entry rate of the total CD8^+^ T cell population). **B – E**; Analyses of the IV^−^ compartment from the mice shown in **Figure 3. B**. Total cell count of IV^−^ CD45^+^ cells normalized per unit tumor volume at the indicated harvest time points. Gated on live, CD45^+^, 24-hr IV^−^ 3-min IV^−^. In **C – E**, cells are gated on live, CD45^+^, Thy1.2^+^, CD8^+^, 24-hr IV^−^ 3-min IV^−^. **C**. Total cell count of IV^−^ CD8^+^ T cells normalized per unit tumor volume. **D**. Total cell count of granzyme B^+^ IV^−^ CD8^+^ TILs normalized to tumor volume. **E**. Total cell count of Ki-67^+^ IV^−^ CD8^+^ TILs normalized to tumor volume. Data shown are a combination of two independent experiments with 7-8 mice per treatment group per experiment and are representative of 2-4 independent experiments. Statistics calculated using an unpaired Student’s t test at each individual time point. In the Figure, ns indicates p>0.05, * indicates p<0.05, ** and indicates p<0.01.

**Figure S4.**
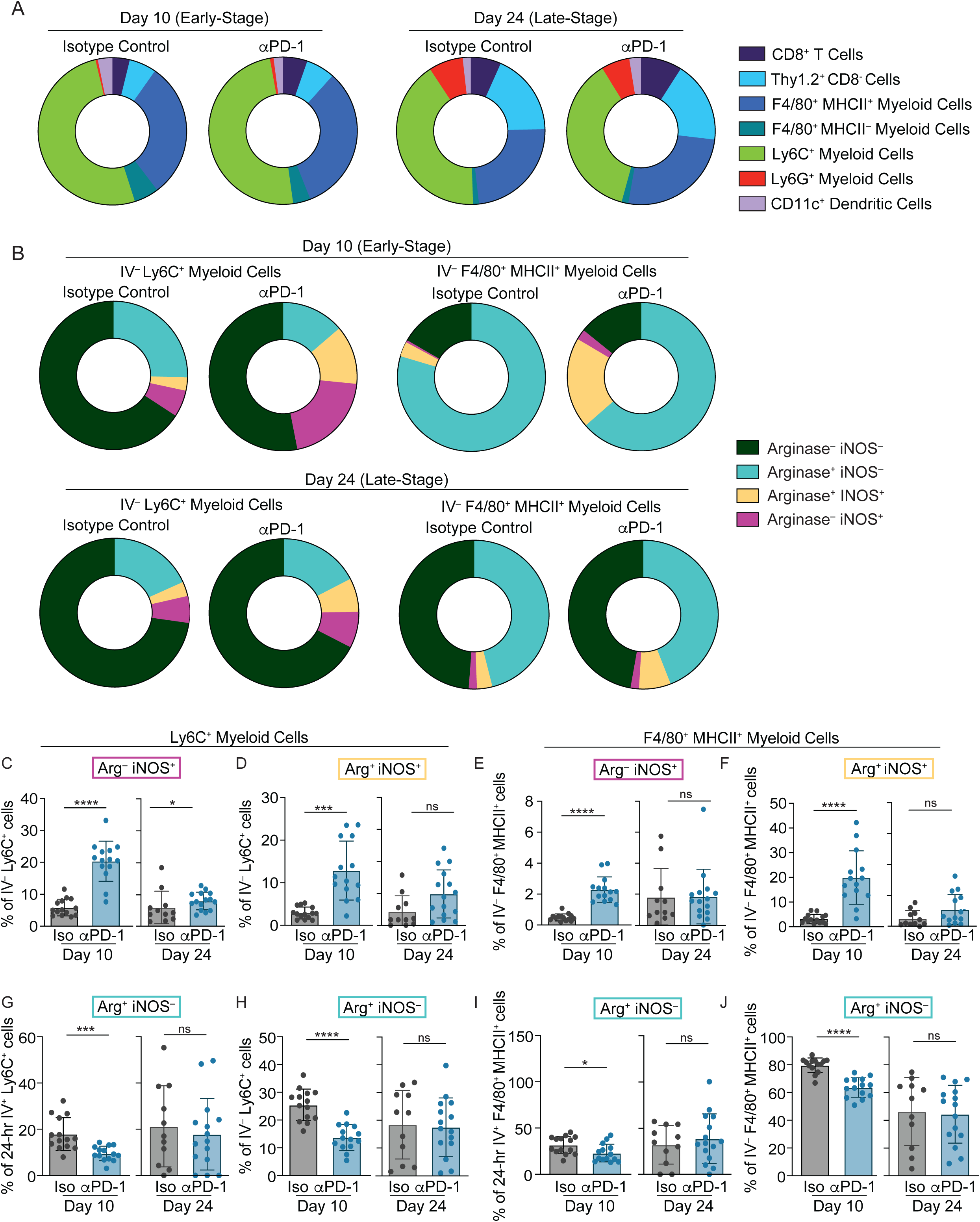
Impact of PD-1 ICI on F4/80^+^ and Ly6C^+^ myeloid cells in the IV^−^ compartment following early vs. late treatment in MC38 tumors. Analyses shown are providing further detail on the mice shown in **Figure 4**. A. Composition of each indicated immune cell lineage within the recently recruited (24-hr IV^+^ 3-min IV^−^) population at the indicated time points (e.g., gating on IV antibody first and then lineage second). Individual lineages further defined as: CD8^+^ T cells: Thy1.2^+^ CD8^+^; Thy1.2^+^ CD8^−^ cells: Thy1.2^+^ CD8^−^; F4/80^+^ MHCII^+^ myeloid cells: F4/80^+^, Ly6G^−^, Ly6C^−^, MHCII^+^, CD11b^+^; F4/80^+^ MHCII^−^ myeloid cells: F4/80^+^, Ly6G^−^, Ly6C^−^, MHCII^−^, CD11b^+^; Ly6C^+^ myeloid cells: F4/80^+^, Ly6G^−^, Ly6C^+^; CD11c^+^ dendritic cells (DCs): F4/80^−^, Ly6G^−^, CD11c^+^, MHCII^+^; Ly6G^+^ myeloid cells: F4/80^−^, Ly6G^+^. In **B – F**, cells are gated on live, CD45^+^, 24-hr IV^−^ 3-min IV^−^ to show the changes in the IV^−^ population. **B**. Composition of IV^−^ F4/80^+^ MHCII^+^ myeloid cells and Ly6C^+^ myeloid cells that express arginase and/or iNOS at the indicated time points. Frequency of the IV^−^ Ly6C^+^ myeloid population expressing iNOS alone (**C**) or iNOS and arginase (**D**). Frequency of the IV^−^ F4/80^+^ MHCII^+^ myeloid population expressing iNOS alone (**E**) or iNOS and arginase (**F**). **G – I** show expression the frequency of myeloid cells expressing arginase but not iNOS (arginase^+^ iNOS^−^). **G – H** show Ly6C^+^ myeloid populations, with **G** showing the frequency of recently recruited (24-hr IV^+^ 3-min IV^+^) and **H** showing unlabeled (24-hr IV^−^ 3-min IV^−^) cells. **I – J** show F4/80^+^ MHCII^+^ myeloid populations, with **I** showing the frequency of recently recruited (24-hr IV^+^ 3-min IV^+^) and **J** showing unlabeled (24-hr IV^−^ 3-min IV^−^) cells. Data shown are a combination of two independent experiments with 7-8 mice treatment group per experiment and are representative of 2-4 independent experiments. Statistical analyses in **Fig. S4C-S4J** were performed using unpaired Student’s t tests. In the Figure, ns indicates p>0.05, * indicates p<0.05, *** indicates p<0.001, and **** indicates p<0.0001.

**Figure S5:**
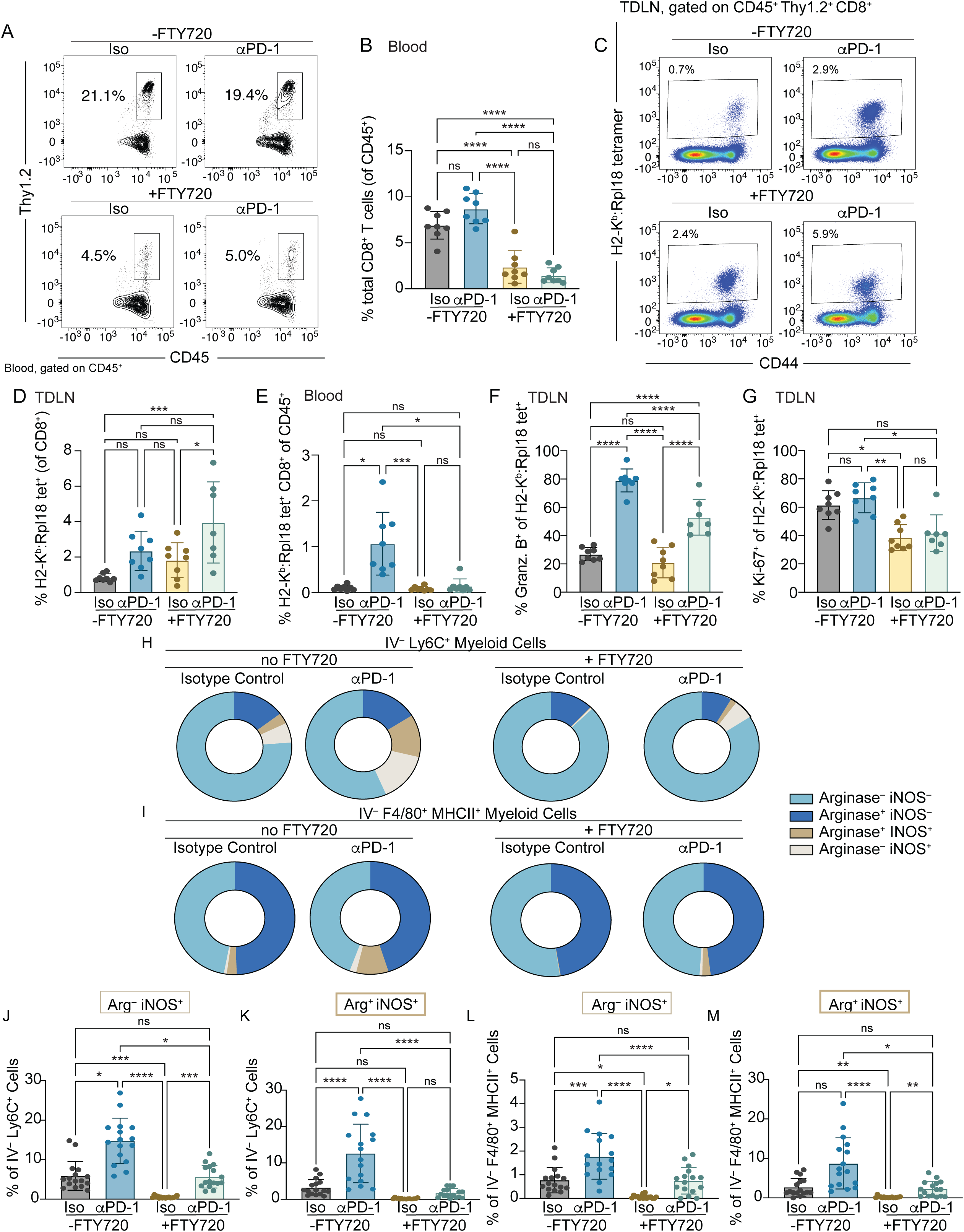
Reducing CD8^+^ T cell trafficking to tumors impairs the protective effects of anti-PD-1 on tumor growth and reduces myeloid cell induction of iNOS. Further analysis of the mice shown in **Figure 5**. A. Representative flow plot of Thy1.2^+^ cells in the blood. Cells are gated on live, CD45^+^ cells. B. Frequency of CD8^+^ T cells of total CD45^+^ cells in the blood. Cells are gated on live, CD45^+^, Thy1.2^+^, CD8^+^. C. Representative flow plots showing H2-K^b^:Rpl18 (KILTFDRL) tetramer (Y axis) vs. CD44 (X axis) in the tumor draining lymph node (TDLN). Cells are gated on live, CD45^+^, Thy1.2^+^, CD8^+^. D. Frequency of H2-K^b^:Rpl18 tetramer-binding cells within the CD8^+^ T cell population in the TDLN. E. Frequency of H2-K^b^:Rpl18 tetramer-binding CD8^+^ T cells of the CD45^+^ population in the blood. Frequency of Granzyme B^+^ cells (F) or Ki-67^+^ cells (G) within the H2-K^b^:Rpl18 tetramer^+^ CD8^+^ T cell population in the TDLN. Data shown in panels A – G are one independent experiment with 7-8 mice per group. Data are representative of 2 independent experiments with 7-8 mice per group. H – I. Composition of IV^−^ Ly6C^+^ myeloid cell and F4/80^+^ MHCII^+^ myeloid cell populations in the tumor that express arginase and/or iNOS. F4/80^+^ MHCII^+^ myeloid cells are gated as F4/80^+^, Ly6C^−^, Ly6G^−^, MHCII^+^, CD11b^+^, while Ly6C^+^ myeloid cells gated as F4/80^+^, Ly6C^+^, Ly6G^−^. Frequency of IV^−^ Ly6C^+^ myeloid cells expressing only iNOS (J) or iNOS and arginase (K). Frequency of IV^−^ F4/80^+^ MHCII^+^ myeloid cells expressing only iNOS (L) or iNOS and arginase (M). Data shown in panels H – M are a combination of two independent experiments with 8 mice per treatment group per experiment. Statistical analyses in Fig. S5B, S5D-S5G and Fig S5J-S5M were performed using One-way ANOVA with multiple comparisons tests. In the Figure, ns indicates p>0.05, * indicates p<0.05, ** indicates p<0.01, *** indicates p<0.001, and **** indicates p<0.0001.

**Figure S6:**
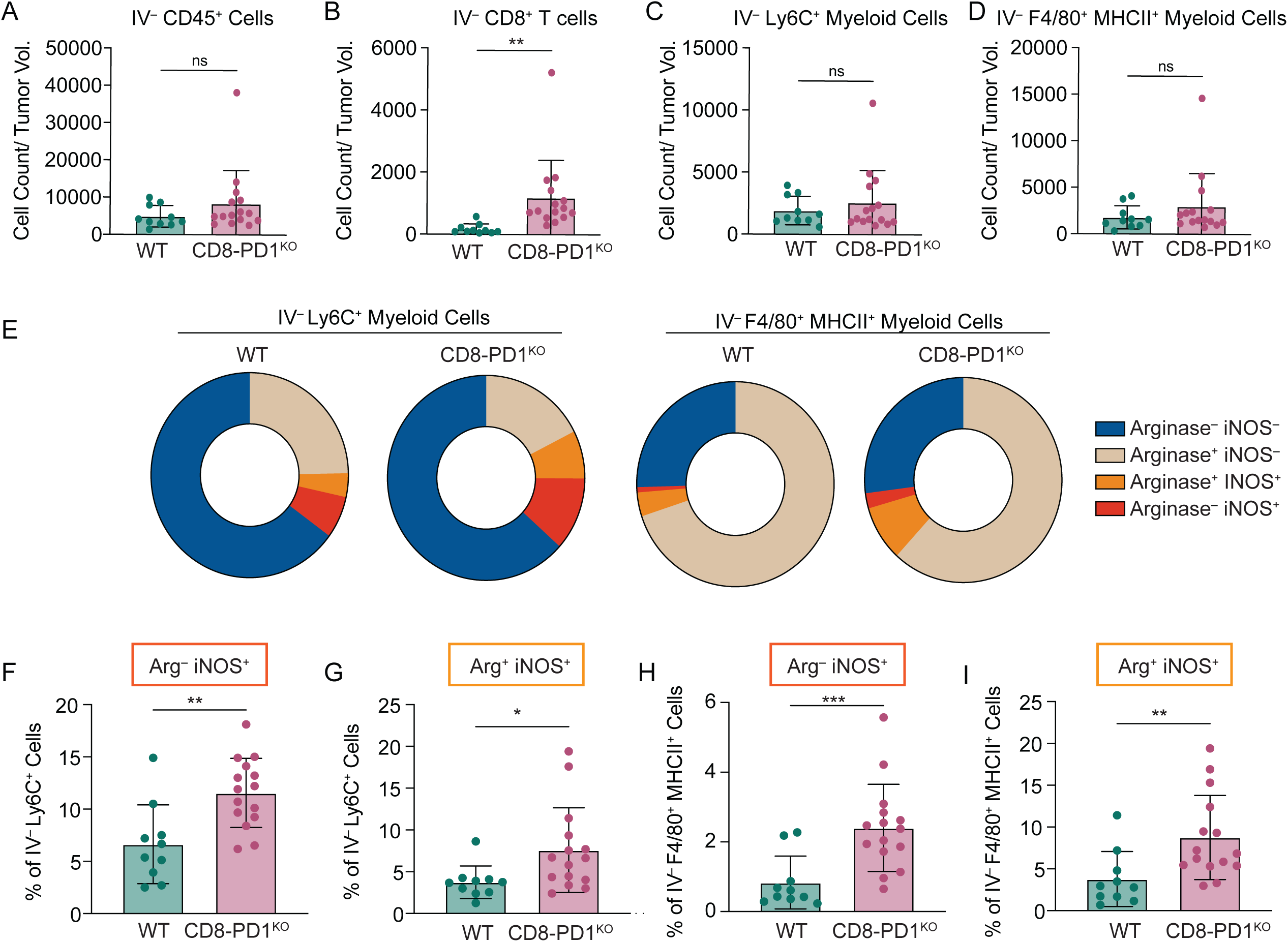
PD-1 deletion in CD8^+^ T cells is sufficient to increase inflammatory myeloid recruitment and create an immunostimulatory TME. Further analysis of the mice shown in **Figure 6**. All cells are minimally gated on live, CD45^+^, and 24-hr IV^−^ 3-min IV^−^ to characterize the unlabeled (IV^−^) compartment. A. Total cell count of IV^−^ CD45^+^ cells normalized per unit tumor volume. B. Total cell count of IV^−^ CD8^+^ T cells normalized per unit tumor volume. Cells further subgated on Thy1.2^+^ CD8^+^. C. Total cell count of IV^−^ Ly6C^+^ cells normalized per unit tumor volume. Cells further subgated on F4/80^+^, Ly6C^+^, Ly6G^−^. D. Total cell count of IV^−^ F4/80^+^ MHCII^+^ cells normalized per unit tumor volume. Cells further subgated on F4/80^+^, Ly6C^−^, Ly6G^−^, MHCII^+^, CD11b^+^. E. Frequency of IV^−^ Ly6C^+^ myeloid cells and F4/80^+^ MHCII^+^ myeloid cells that express arginase and/or iNOS. Frequency of IV^−^ Ly6C^+^ myeloid cells expressing only iNOS (F) or iNOS and arginase (G). Frequency of IV^−^ F4/80^+^ MHCII^+^ myeloid cells expressing only iNOS (H) or iNOS and arginase (I). Data shown in all panels are a combination of two independent experiments with 4-7 mice per treatment group per experiment. Statistical analyses in Fig. S6A-5D and S6F-6I were performed using unpaired Student’s t tests. In the Figure, ns indicates p>0.05, * indicates p<0.05, ** indicates p<0.01, and *** indicates p<0.001.

